# DRAXIN regulates interhemispheric fissure remodelling to influence the extent of corpus callosum formation

**DOI:** 10.1101/2020.07.29.227827

**Authors:** Laura Morcom, Timothy J Edwards, Eric Rider, Dorothy Jones-Davis, Jonathan WC Lim, Kok-Siong Chen, Ryan Dean, Jens Bunt, Yunan Ye, llan Gobius, Rodrigo Suárez, Simone Mandelstam, Elliott H Sherr, Linda J Richards

## Abstract

Corpus callosum dysgenesis (CCD) is a congenital disorder that incorporates either partial or complete absence of the largest cerebral commissure. Remodelling of the interhemispheric fissure (IHF) provides a substrate for callosal axons to cross between hemispheres, and its failure is the main cause of complete CCD. However, it is unclear whether defects in this process could give rise to the heterogeneity of expressivity and phenotypes seen in human cases of CCD. We identify incomplete IHF remodelling as the key structural correlate for the range of callosal abnormalities in inbred and outcrossed BTBR mouse strains, as well as in humans with partial CCD. We identify an eight base pair deletion in *Draxin* and misregulated astroglial and leptomeningeal proliferation as genetic and cellular factors for variable IHF remodelling and CCD in BTBR acallosal strains. These findings support a model where genetic events determine corpus callosum structure by influencing leptomeningeal-astroglial interactions at the IHF.

## Introduction

The corpus callosum (CC) is the largest white matter tract that mediates information transfer between brain hemispheres in placental mammals. In humans, it fails to form normally in approximately 1:4000 live births resulting in a group of conditions collectively termed CC dysgenesis (CCD; Glass et al., 2008). CCD incorporates complete and partial congenital absence of the CC, as well as hypo- and hyperplasia (thinning or thickening, respectively) of the CC. Each of these structural phenotypes can variably impact brain function and organization (Brown and Paul, 2019; Edwards et al., 2014; Paul et al., 2007), but the precise developmental mechanisms that could give rise to these diverse CCD phenotypes remain unknown.

CC formation is dependent on a prior sequence of developmental processes: cellular proliferation, migration, axonal elongation, guidance and targeting (Donahoo and Richards, 2009; Edwards et al., 2014; Morcom et al., 2015). Callosal axons derived from cells within the cingulate and neocortices elongate and cross the telencephalic midline in a region of the septum termed the commissural plate (Moldrich et al., 2010; Rakic and Yakovlev, 1968). We previously demonstrated that in order for callosal axons to cross the midline, the interhemispheric fissure (IHF) that separates the telencephalic hemispheres must be remodelled to form a permissive substrate (Gobius et al., 2016). IHF remodelling is mediated by intercalation of specialized astroglia, known as the midline zipper glia (MZG), across the IHF (Silver et al., 1993). This process does not occur in naturally acallosal mammalian marsupial and monotreme species (Gobius et al., 2017) and its failure appears to be a major cause of complete CCD in humans (Gobius et al., 2016). When IHF remodelling does not proceed normally in placental mammals, callosal axons do not cross the midline and can instead form longitudinal tracts in the ipsilateral hemisphere that are known as Probst bundles (Probst, 1901). Although IHF remodelling is a prerequisite for CC formation, our understanding of the cellular and genetic factors involved is incomplete. Moreover, it is unknown whether disruptions to this process could account for the spectrum of commissural phenotypes seen in CCD, or if these are due to independent developmental mechanisms.

Absence or dysgenesis of the hippocampal commissure (HC) is frequently observed to co-occur with CCD in humans, indicating that these commissures rely on common developmental programs (Hetts et al., 2006). Analogously, the BTBR T+ Itpr3tf/J (BTBR) mouse has a severe commissural phenotype incorporating complete CCD and HC dysgenesis (Wahlsten, 2003). Using the F2 generation of BTBR x C57Bl/6J (C57) intercross, which displays variable CCD and HC dysgenesis, we previously demonstrated that a highly penetrant locus on chromosome 4 was associated with CC and HC size (Jones-Davis et al., 2013). This suggests that the degree of dysgenesis of two major telencephalic commissures may result from disruption of a single developmental process.

Here, we investigate the underlying genetic and developmental mechanisms leading to diverse CCD phenotypes in BTBR mice and the BTBR x C57 cross. CCD severity and the extent of HC dysgenesis in these mouse strains was strongly associated with abnormal retention of the IHF, and thus the degree to which IHF remodelling was incomplete. Moreover, we describe an eight base pair deletion in the axon guidance gene *Draxin,* which truncates and ablates normal DRAXIN protein expression in BTBR mice. Inheritance of the *Draxin* mutation is a driver of defective IHF remodelling and subsequent CCD and HC dysgenesis in BTBR and BTBR x C57 mice. Mis-regulated cellular proliferation of MZG and leptomeningeal cells were both identified as the cellular correlates for failed MZG-mediated IHF remodelling and interhemispheric tract formation in BTBR mice. Finally, we identify incomplete IHF remodelling in a cohort of human individuals with partial CCD. Together our results suggest diverse CCD phenotypes can arise from a single genetic event that variably disrupts IHF remodelling and consequently the amount of substrate available for CC and HC axons to cross the midline and therefore provides the first aetiology associated with partial CCD.

## Materials and methods

### Animals

BTBR, CD1, and C57 mice were bred and tested at The University of Queensland according to the Australian Code of Practice for the Care and Use of Animals for Scientific Purposes and with prior ethics approval from The University of Queensland Animal Ethics Committee. To generate BTBR x C57 N2 mice with varying degrees of CCD, an N1 intercross was produced, and N1 females were crossed with BTBR males to generate N2 mice for analysis. The day of birth was designated postnatal day (P)0. Time-mated females were obtained by housing male and female mice together overnight and the following morning was designated embryonic day (E)0. Animals were anaesthetised and collected as previously described (Suárez et al., 2014). For EdU labelling, pregnant dams were given an intraperitoneal injection of 5-ethynyl-2′-deoxyuridine (EdU; 5 mg per kg body weight at E14 or 7.5 mg per kg body weight at E12 and E13, Invitrogen) and embryos were collected 24 hours later. Sex was not determined for embryonic studies. A total of 112 adult BTBR x C57 N2 mice were analysed (n = 47 males, n = 65 females). For the analysis of adult mouse tissue, mice were perfused at 12 weeks of age.

Genotyping of BTBR x C57 N2 mice was performed by PCR of single nucleotide polymorphisms (SNPs) associated with CC size in the BTBR x C57 F2 intercross as previously described by us (Jones-Davis et al., 2013). Amplicons were generated for SNP regions using the following primers: Chromosome 4, SNP rs6397070, forward TTTATGGCTGGGGACTTCAG and reverse CGAATCCAAAGCTCTCTTGC; Chromosome 9, SNP rs29890894, forward AGCTTGGTGGCATCCATATC, reverse GCACTCTCCCTACTGCTTGG; Chromosome 15, SNP rs31781085, forward GATCGTTGCAGTGACCACAC, reverse GCTGATTGGCAGGTTCTGAT. Genotyping for the *Draxin* mutation was performed by PCR using the following primers: wildtype forward AGACGGTCCCTGCGTCTC, mutant forward GTCGCAGACGGTCCCTTG, and common reverse AGGCTTCCCAGATGACACTC. Sanger sequencing was performed at the Australian Genome Research Facility.

### Human participants

Participants were enrolled in an ongoing study at The University of Queensland from October 2014 with approval from The University of Queensland Research Ethics committee. Ten individuals with partial CCD aged 21-72 (mean age = 40 years, standard deviation = 17.73) and nine neurotypical individuals (mean age = 34.89, standard deviation = 16.77) provided written informed consent. MRI scans of all participants were reviewed by a neuroradiologist (S.M.), who has extensive experience in human brain malformations. The groups did not differ significantly in either age or gender. Genetic information to determine cause of CCD was not collected for this study.

### MRI scans

MRI scanning was performed using the same parameters and cohort of BTBR x C57 N2 adult mice as described in Edwards et al. (2020). Briefly, skulls were post-fixed in 4% paraformaldehyde for at least 48 hours after perfusion. Following this, each BTBR x C57 N2 adult mouse skull (with brain *in* situ) was washed in phosphate buffered saline (PBS) with 0.2% sodium azide. Skulls were immersed in Fomblin Y-LVAC fluid (Solvay Solexis, Bollate, Italy) and air was actively removed from each sample via vacuum pumping prior to MRI scanning. All skulls (n = 112) initially underwent two-dimensional scans measuring diffusion in the mediolateral direction to identify the commissural phenotype. From this analysis, a subset of brains of each of complete CCD (n = 10), partial CCD (n = 11) and full CC (n = 10) phenotypes were dissected from skulls, incubated for four days in 0.2% gadopentetate dimeglumine (Magnevist, Berlex Imaging, Wayne, NJ, USA), and scanned with a 16.4 Tesla Bruker Avance MRI scanner using a FLASH sequence that was described previously (Schanze et al., 2018): FLASH sequence (voxel size = 0.03 × 0.03 × 0.03 mm, MTX 654 × 380 × 280, FOV 19.6 × 11.4 × 8.4 mm, TR = 50 ms, TE = 12 ms, flip angle of 30°). Three adult BTBR inbred mouse brains were scanned using identical processing steps. C57 mouse brain scans were acquired previously for Ullmann and colleagues (2013), and were kindly provided by Dr. Nyoman Kurniawan (Centre for Advanced Imaging, The University of Queensland, Australia).

Human participants underwent MRI at the Centre for Advanced Imaging (The University of Queensland) using a 7 Tesla Siemens Magnetom whole body MRI scanner. Structural MRI data was acquired as described previously (Hearne et al., 2019): at 7T, MP2RAGE sequence (voxel size = 0.75 × 0.75 × 0.75 mm, MTX 256 × 300 × 320, FOV 192 × 225 × 240 mm, TR = 4300 ms, TE = 3.44 ms, TI = 840 / 2370 ms, flip angle of 5°, FOV 192 × 225 × 240 mm).

### MRI anatomical measurements

Commissure and brain sizes were measured in OsiriX (v 5.8.5; Rosset et al., 2004), blind to animal genotype. All commissure lengths and areas were measured in the midsagittal plane on single direction diffusion scans. Anteroposterior CC length was measured using the straight length tool from the most anterior point to the most posterior point of the CC. HC length was measured using the straight length tool from the dorsal-most aspect of the HC (inferior to the CC) to the ventral-most aspect of the HC. HC area was measured using the closed polygon tool using the same superior and inferior boundaries for HC length. Anterior commissure area was measured using the closed polygon tool. Brain length was measured using the straight length tool, from the anterior aspect of the frontal pole to the posterior-most aspect of the cerebellum. IHF and CC measurements from human MRI were measured from representative images displayed within figures using ITK-SNAP v3.8.0 (Yushkevich et al., 2006).

### Immunohistochemistry and tissue staining

Immunohistochemistry was performed on 50 μm tissue sections as previously described (Moldrich et al., 2010). Primary antibodies used: Sheep anti-DRAXIN (1:250; AF6149, R&D systems), mouse anti-human KI67 (1:500; 550609, BD Pharmingen), mouse anti-GAP43 (1:500; AB1987, Millipore), rabbit anti-GFAP (1:500; Z0334, Dako), mouse anti-GLAST (or EAAT1; 1:500; ab49643, Abcam), rabbit anti-GLAST (or EAAT1; 1:250; ab416, Abcam), chicken anti-LAMININ (1:250; LS-C96142, LSBio), rabbit anti-LAMININ (1:250; L9393, Sigma), rat anti-NESTIN (AB 2235915, DSHB), and rabbit anti-SOX9 (1:500, AB553, Merck). Secondary antibodies were Alexa Fluor IgG antibodies (1:500, Invitrogen) or biotinylated IgG antibodies (1:500 or 1:1000, Jackson Laboratories) used in conjunction with Alexa Fluor 647-conjugated Streptavidin (1:500, Invitrogen) amplification. EdU labelling was performed using the Click-iT EdU Alexa Fluor 488 or Alexa Fluor 555 Imaging kits (Invitrogen) according to the manufacturer’s instructions. Cell nuclei were labelled using 4’,6-diamidino-2-phenylindole dihydrochloride (DAPI, Invitrogen) and coverslipped using ProLong Gold anti-fade reagent (Invitrogen) as mounting media.

### In situ hybridization

*In situ* hybridization was performed as previously described (Moldrich et al., 2010), with the following minor modifications: Fast red (Roche) was applied to detect probes with fluorescence. For fluorescent *in situ* hybridization against *Draxin* mRNA in wildtype CD1 mice, the *Draxin* CDS was amplified by PCR using the following primer pairs from the Allen Brain Atlas (Lein et al., 2007): CAGGGAGGTTTAGGACAAACAG and TGTAGGAGCTGAGGGAAAGAAG. The *Draxin* CDS was subsequently cloned into the pGEM-T Vector (Promega USA) and sequences were verified. Digoxygenin-labelled (DIG RNA labelling mix; Roche) antisense riboprobes were also generated in a similar manner from the *Draxin* CDS amplified from BTBR and C57 E15 telencephalic midline tissue using the following primer pairs: CGACAGGGAGAGCCAATG and GTACTGGGCGTACACCTGCT.

### Generation of BTBR and C57 Draxin expression plasmids

The *Draxin* CDS was amplified from BTBR and C57 E15 telencephalic midline tissue using the following primer pairs containing added EcoRI and NotI restriction sites: GAATTCGACAGGGAGAGCCAATG and GCGGCCGCGTACTGGGCGTACACCTGCT. The amplified sequence was digested with EcoRI and NotI and subsequently inserted into the pPBCAG-IRES-GFP (pPBCAGIG) expression plasmid (Chen et al., 2020) via the corresponding restriction sites. Successful cloning of the *Draxin* CDS into the pPBCAGIG plasmid was verified by Sanger sequencing.

### Western blot

Whole-cell protein extracts were prepared from transfected HEK293T cells and dissected midline tissue as described previously (Bunt et al., 2010). Protein extracts were cleared by centrifugation and used for Western blotting as described previously (Bunt et al., 2017). Primary antibodies used for immunoblotting were sheep anti-DRAXIN (AF6149, R&D systems, 1 ug/mL), goat anti-β-ACTIN (AB0145-200, SICGEN, 1:1000 or 3 ug/mL) and rabbit anti-GFP (A-6455, Invitrogen, 1:1000). The secondary antibodies used were Sheep IgG (H&L) antibody DyLight 800 conjugated (Rockland Immunochemicals Inc., 1:15000), IRDye 680LT donkey anti-rabbit (LI-COR, 1:15000) and IRDye 800CW donkey anti-goat (LI-COR, 1:15000). Immunoblotted membranes were imaged using the Odyssey Classic (LI-COR) and Image Studio 5 software (LI-COR).

### Image acquisition and analysis

Microscopy for fluorescence immunohistochemistry or *in situ* hybridization was performed using either an inverted Zeiss Axio-Observer fitted with a W1 Yokogawa spinning disk module, Hamamatsu Flash4.0 sCMOS camera and Slidebook 6 software or an inverted Nikon TiE fitted with a Spectral Applied Research Diskovery spinning disk module, Hamamatsu Flash4.0 sCMOS camera and Nikon NIS software. Pseudocoloured image projections of ~10 – 20μm thick z-stacks were acquired. Images of chromogenic *in situ* hybridization samples were acquired on a Zeiss upright Axio-Imager Z1 microscope with Axio-Cam HRm camera and Zen software (Carl Zeiss). Images were cropped, sized, and contrast-brightness enhanced for presentation with ImageJ and Photoshop software (Adobe).

The ratio of IHF length to total telencephalic midline length was quantified in ImageJ freeware (National Institutes of Health, Bethesda, USA) as previously described (Gobius et al., 2016). The IHF width was measured from LAMININ and GLAST stained tissue sections in two different regions: 1) ~ 5 μm from the base of the IHF and 2) at the most rostral region where GLAST-positive MZG fibres attach to the IHF surface which coincided with the corticoseptal boundary. The IHF width at the base (region 1) was then expressed as a ratio over the IHF width at the corticoseptal boundary (region 2) in order to reflect changes in the compression of the IHF at the base prior to IHF remodelling. Fluorescence intensity of GLAST-positive and NESTIN-positive MZG fibres in a region of interest ~100 × 200 μm (medial-lateral x rostral-caudal) along the IHF surface was measured in ImageJ v1.52i freeware from multiple intensity projection 2D images generated from 3D z-stacks. The number of SOX9-positive MZG cell bodies was counted manually using the Cell Counter plugin in ImageJ v1.51s freeware from a region of interest at the surface of the IHF measuring ~10 × 200 μm (medial-lateral x rostral-caudal). For cell proliferation assays, the telencephalic hinge was outlined as a region of interest in Imaris from 3D z-stacks. The number of DAPI-positive, EdU-positive, KI67-positive cells were automatically counted using the Imaris spots function as previously described (Faridar et al., 2014). The colocalization function was used to determine co-labelled spots within 4 μm (MZG) or 5 μm (leptomeninges) of each other. All counts were performed blind to the experimental conditions and normalised to the number of DAPI-positive cells or the volume of the region of interest.

### Experimental design and statistical analysis

Statistical tests were performed in GraphPad Prism (v.7 and v.8), and p < 0.05 was considered significant. Statistical testing was performed for all quantitative analyses – the relevant statistical test performed, number of biological replicates, and level of significance is described for all quantitative results in figures or their legends, and in Supplementary Table 1. For qualitative comparisons (figures 4 and 8), a minimum of three animals were used for each experiment.

## Results

### The CC and HC are variably malformed in BTBR x C57 N2 mice

We previously demonstrated that BTBR x C57 N2 littermates display a spectrum of CCD phenotypes (Jones-Davis et al, 2013; Edwards et al., 2020). Here, we further classified these phenotypes into full CC (CC length ≥ 3 mm, according to typical C57 wildtype CC lengths in Jones-Davis et al., 2013; Figure 1A-B), partial CCD (CC length > 0 and < 3 mm), and complete CCD (CC length = 0 mm; Figure 1A-B). We identified variable HC size in animals with complete CCD, suggesting that distinct subpopulations of CCD with variable HC dysgenesis may occur within the BTBR x C57 N2 mouse (Figure 1C-D). Furthermore, the severity of CCD was correlated with HC dysgenesis; HC length was reduced in partial CCD and complete CCD compared to full CC littermates, and was reduced in complete CCD compared to partial CCD (Figure 1C and Table S1). These results demonstrate that the BTBR x C57 N2 mice display a range of CC phenotypes suitable for further investigation of underlying aetiologies. Moreover, the association between CC length and HC length (as observed in our initial QTL-based manuscript on CC and HC morphology; Jones-Davis et al., 2013), suggests that a common aetiology may underlie the observed variance in commissural size in this mouse cross.

**Figure 1:**
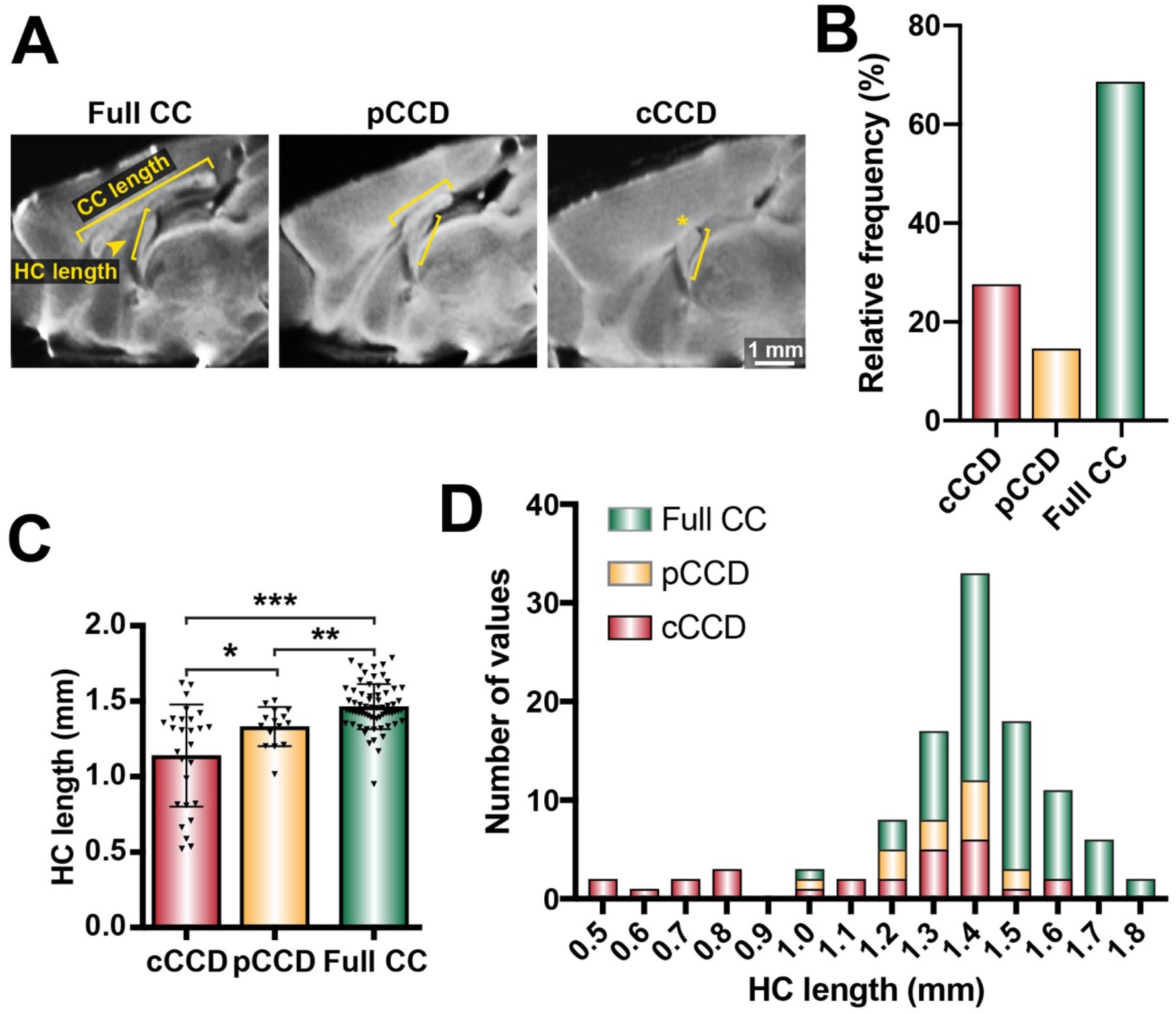
Distribution of commissural size in BTBR x C57 N2 mice. (A) CC length and HC length were measured on single diffusion direction MRI scans for n = 112 adult BTBR x C57 N2 mice. (B) The relative frequency of three distinct subsets of CC phenotypes based on CC length: complete CCD (cCCD; red), partial CCD (pCCD; yellow), and normal CC length (Full CC; green). (C) Group-wise comparison between callosal phenotypes for HC length. (D) Stacked histogram of HC length for each callosal phenotype. Data represented as mean ± SEM. Welch’s t-test: * p < 0.05; ** p < 0.01; *** p < 0.001.

### CC and HC malformations are associated with defects in IHF remodelling in the BTBR x C57 N2 and BTBR parental mouse strain

To investigate potential structural correlates of complete and partial CCD, we first examined the midline of BTBR and BTBR x C57 N2 mice. We previously demonstrated that remodelling of the IHF is a critical developmental step required for subsequent midline crossing of callosal axons, and that failure of this process to occur results in an unfused septum; an MRI feature that is strongly associated with complete CCD in humans (Gobius et al., 2016). To determine whether incomplete IHF remodelling may underlie the spectrum of CCD seen in the BTBR x C57 N2 mouse, we acquired high resolution structural MRI scans of adult C57 and complete CCD BTBR mice, as well of a subset of BTBR x C57 N2 mice with complete and partial CCD.

C57 adult mice demonstrated a fused septum and IHF positioned anteriorly and superiorly to the CC and HC (Figure 2A). In contrast, the acallosal BTBR adult mouse demonstrated an unfused septum and an IHF that is aberrantly retained across almost the full extent of the telencephalic midline (Figure 2B, yellow bracket). Consistent with what has been shown previously, the only crossing white matter identified in the dorsal telencephalon of the BTBR mouse was a small, ventrally positioned HC (Wahlsten et al., 2003). Full CC BTBR x C57 N2 mice demonstrated similar midline anatomy to the C57 mice (Figure 2C). Partial CCD mice demonstrated an intact HC; however, the anterior-posterior extent of the CC was reduced, and was associated with dorsal retention of the IHF, suggesting that the IHF had not been fully remodelled (Figure 2D). In support of an aetiologic link between an IHF remodelling defect and CCD severity, complete CCD mice had a more severe IHF remodelling defect, with near complete retention of the IHF (Figure 2E). A subset of complete CCD mice displayed a more severe IHF phenotype with an associated dysgenesis of the HC that recapitulates that seen in the BTBR parental strain (left panels, Figure 2E), compared to complete CCD mice with an intact HC (right panels, Figure 2E). Together, these data suggest that the underlying pathogenesis of CCD in the BTBR x C57 N2 mouse is incomplete IHF remodelling leading to aberrant retention of the IHF. This further suggests that the severity of the HC and callosal phenotypes are related to the degree of IHF remodelling that occurs.

**Figure 2:**
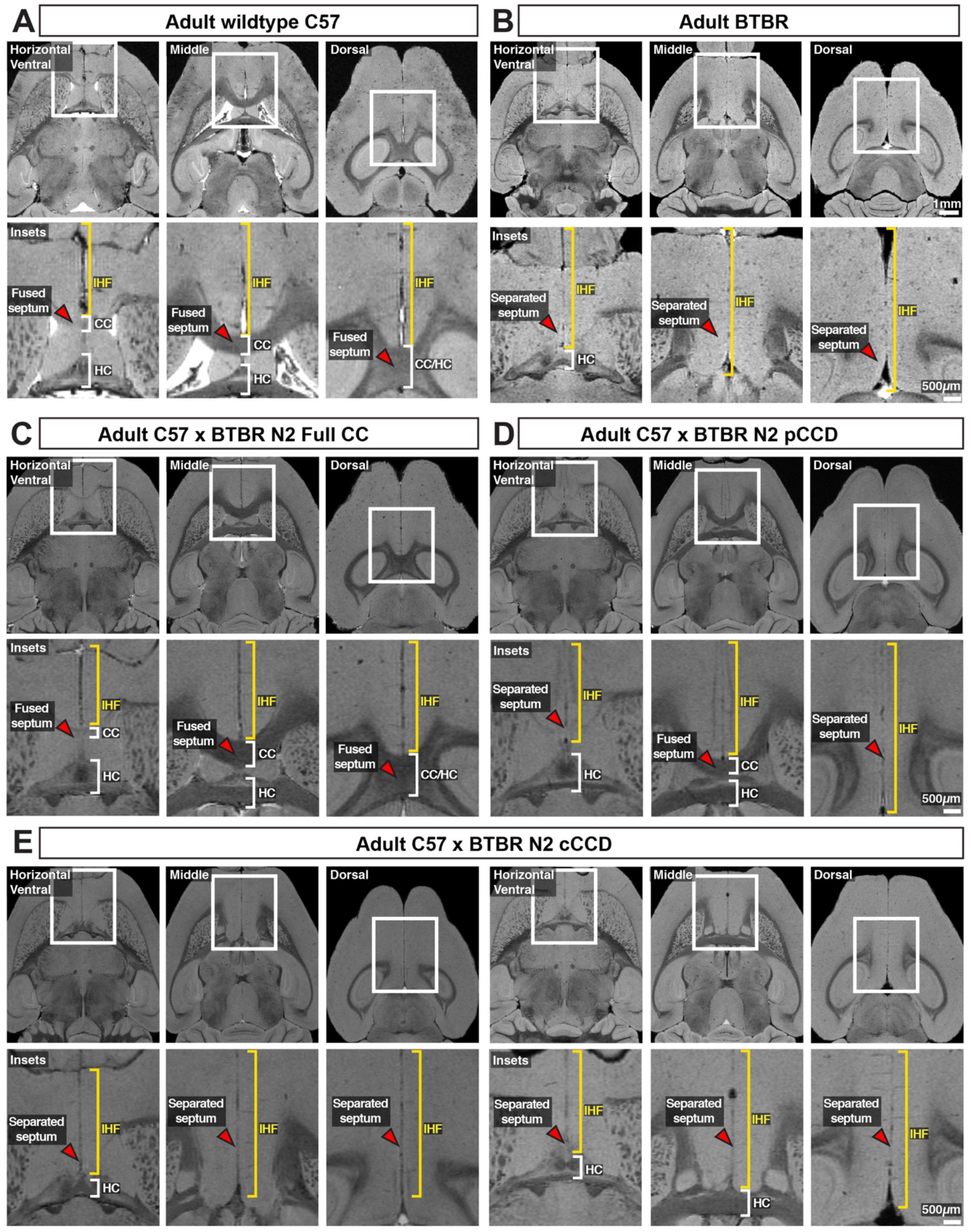
Severity of CC and HC dysgenesis is associated with the extent of retention of the IHF in BTBR x C57 N2 mice. Axial structural MRI slices and insets (white rectangles) of telencephalic midline anatomy in adult wildtype C57 (A) and acallosal BTBR parental mice (B), as well as adult BTBR x C57 N2 mice with distinct commissural phenotypes (C – E). The IHF is indicated with yellow brackets, the CC and HC are indicated with white brackets, and the septum is indicated with red arrowheads. n = 3 for C57 and BTBR parental strains, n = 10 for each callosal condition for the BTBR x C57 N2 cross. cCCD = complete CCD, pCCD = partial CCD.

### Retention of the IHF with an unfused or absent septum is associated with the degree of partial CCD in humans

An unfused septum and deep IHF are invariably present in humans with complete CCD, who commonly have associated HC malformations (Gobius et al., 2016; Hetts et al., 2006). To determine whether partial CCD individuals display similar septal and IHF abnormalities, we examined the septum and IHF in structural MRI scans of 10 adult individuals with partial CCD and compared these to nine neurotypical controls (Figure 3A and Figure S1). Partial CCD individuals demonstrated variably positioned CC remnants comprising one or more, but not all, of the normal corpus callosum segments. These individuals often had other mild brain abnormalities that were deemed to be not related to midline formation and IHF remodelling, except for two individuals that demonstrated interhemispheric cysts.

**Figure 3:**
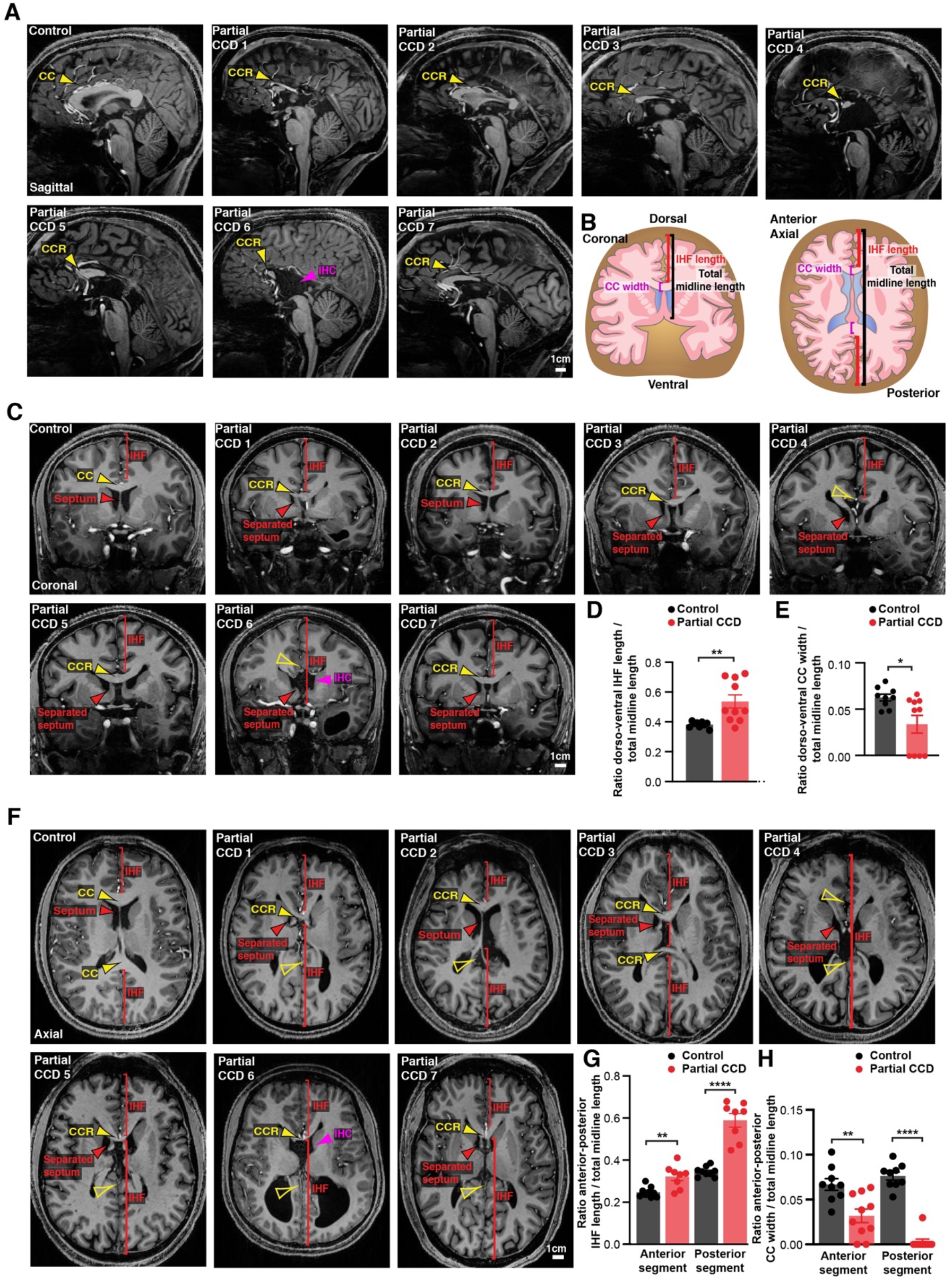
Structural MRI study of IHF and septal defects associated with partial CCD in humans. Representative sagittal (A), coronal (C) and axial (F) slices from T1-weighted structural scans on 9 neurotypical (control) individuals and 10 individuals with partial CCD. The CC or the CC remnant (CCR) is indicated with yellow arrowheads, interhemispheric cysts (IHC) are indicated with magenta arrowheads, the IHF extent is indicated with red brackets and the septum is indicated with red arrowheads. The IHF length and CC width were measured from coronal and axial images and normalised to the length of the midline, as quantified in D, E, G, H and schematised in B. Data is represented as mean ± SEM. * p <0.05; ** p< 0.01, **** p< 0.0001 as determined with unpaired t tests or Mann-Whitney tests. See Figure S1 for further subjects and quantification.

In neurotypical individuals, the CC extends dorsal, anterior and posterior to the septum, which is fully fused except in a minority of individuals that demonstrate cavum septum pellucidum – a normal anatomical variant that forms ventral to the corpus callosum (Schwidde, 1952). Partial CCD individuals demonstrated significantly increased IHF length in the dorso-ventral and anterior-posterior axis (Figure 3B-D and 3F-G, Figure S1B and S1C, and Table S1), and a significant reduction in the length of fused septum along the anterior-posterior axis as normalised to total midline length (Figure S1D, Table S1). All partial CCD individuals displayed increased length of the posterior IHF in the axial plane outside of the range measured in neurotypical individuals (Figure 3G). Increased length of the anterior and posterior IHF indicates a decrease in IHF remodelling and septal fusion, and therefore a reduced amount of tissue available for callosal midline crossing. This correlated with a reduction in CC width in partial CCD; the posterior CC being the most severely affected since it was absent in all but one individual with partial CCD (Figure 3E and 3H). Notably, the CC remnant in partial CCD individuals was often displaced ventrally within the fused septum where it is not normally evident in neurotypical controls (Figures 3C and S1B). Several individuals displayed absence of the septal leaves that occurred in association with an interhemispheric cyst (Partial CCD subject 6 and 10; Figures 3 and S1). These results suggest that partial CCD is associated with incomplete IHF remodelling and septal fusion in our cohort. Therefore, developmental failure of IHF remodelling appears to be consistently associated with a spectrum of CCD phenotypes in mice and humans.

### A deletion within Draxin in the parental BTBR strain is associated with loss of Draxin expression at the telencephalic midline

Families with CCD can exhibit variable expressivity when carrying the same inherited pathogenic gene variant (Marsh et al., 2017). Because complete and partial CCD BTBR x C57 N2 littermates both display an IHF remodelling defect, the severity of which is associated with the severity of CCD, we sought to determine whether they share a genetic aetiology. Our previously described linkage analysis demonstrated a high LOD score QTL for CC and HC anatomy on the distal end of chromosome 4 (Jones-Davis et al., 2013). This locus was found to overlap with an eight-base pair deletion introducing a premature stop codon in exon 2 of *Draxin* in the BTBR strain (Figure 4A-B). *Draxin* is a promising candidate to explain CCD in the BTBR x C57 N2 mouse, since *Draxin* knockout mice display CCD (Ahmed et al., 2011; Islam et al., 2009). Moreover, mutations in the gene encoding the DRAXIN receptor, DCC, are also associated with CCD in mice and humans (Fazeli et al., 1997; Finger et al., 2002; Fothergill et al., 2013; Jamuar et al., 2017; Marsh et al., 2018; Marsh et al., 2017). We generated *in situ* riboprobes for wildtype and mutant *Draxin* using mRNA isolated from C57 and BTBR mice, respectively, and examined the mRNA expression in the BTBR x C57 N2 and the BTBR parental mice. *In situ* hybridization with 3’ *Draxin* antisense probes from both strains revealed that *Draxin* is highly expressed in the cingulate cortex and the septum in mid-horizontal sections of the telencephalic midline of C57 mice, consistent with previous findings (Figure 5D; Islam et al., 2009). *Draxin* mRNA was comparable between C57 mice and mice homozygous for the *Draxin* mutation (BTBR, and BTBR x C57 N2 mice), but protein expression was undetectable in tissue using a polyclonal antibody generated from an immunogen of whole human DRAXIN via immunohistochemistry and western blot (Figure 4B, 4E and Figure S2). However, western blot revealed that expression of the BTBR *Draxin* coding sequence in HEK293T cells produced DRAXIN protein of reduced molecular weight compared to the C57 *Draxin* coding sequence (Figure 4D and Figure S2). These results suggest that the eight base pair deletion in *Draxin* truncates DRAXIN and disrupts normal protein expression *in vivo*.

**Figure 4:**
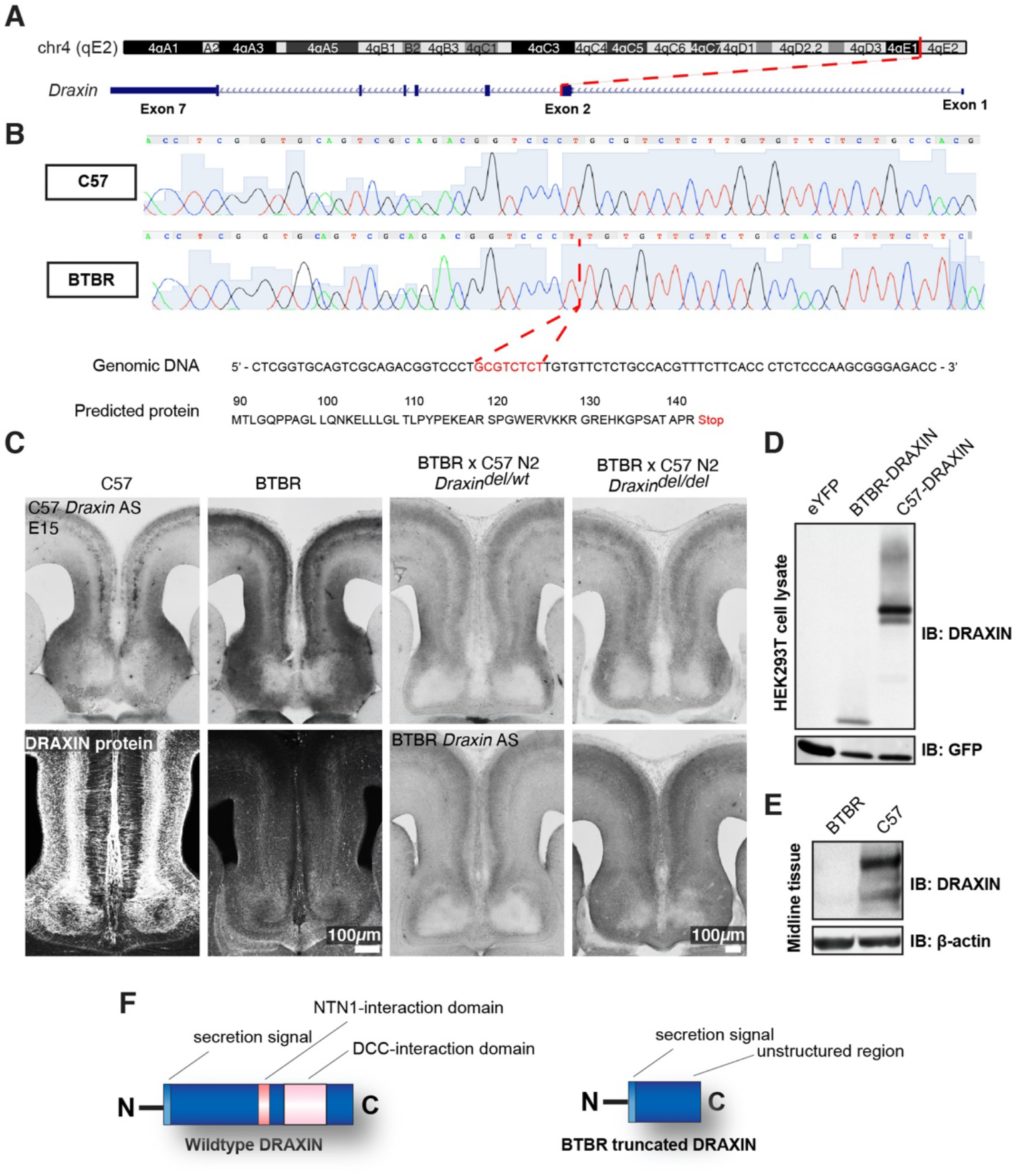
An eight base pair deletion in *Draxin* in BTBR strains truncates and reduces DRAXIN protein expression. Exome sequencing of a candidate region on mouse chromosome 4 (A) and confirmatory Sanger sequencing found an eight base pair deletion in *Draxin*, which introduces a premature stop codon (B), predicted to truncate DRAXIN protein (B, F). (C) *In situ* hybridization against 3’ C57 or BTBR *Draxin* demonstrates a similar pattern of *Draxin* mRNA expression in C57 and BTBR parental strains and in the BTBR x C57 N2 mice. Fluorescent immunohistochemistry for DRAXIN protein (bottom left panels) demonstrates that DRAXIN is highly expressed in C57 mice at E15 but not in BTBR mice. (D) Cell lysates derived from HEK293T cells expressing pCag-eYFP, or pCag-iresGFP with either BTBR or C57 *Draxin* coding sequences were incubated with anti-DRAXIN and anti-GFP antibodies. Specific bands at ~18kD and ~40-45kD are shown and demonstrate that BTBR *Draxin* produces a protein of reduced molecular weight, indicating truncation. (E) Midline tissue lysates from E15 C57 and BTBR mice incubated with anti-DRAXIN and anti-β-ACTIN antibodies reveal specific bands at ~40-60kD and ~42kD respectively, indicating DRAXIN expression is severely reduced in BTBR mice. See related Figure S2.

**Figure 5:**
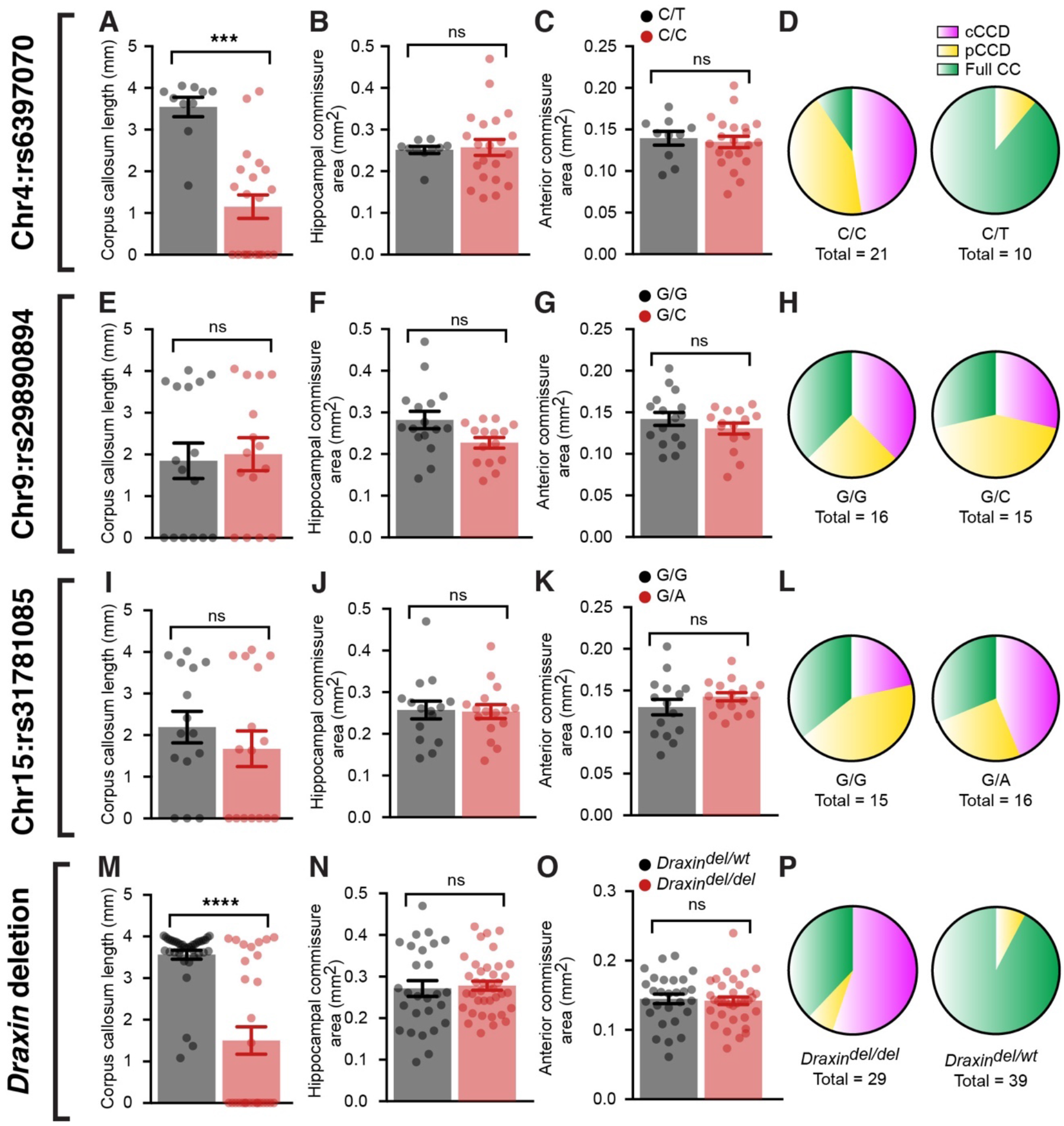
BTBR allele on chromosome 4 and a deletion in *Draxin* are associated with CCD in BTBR x C57 N2 mice. Group-wise comparison for marker SNPs at candidate genetic loci compared with CC length (A, E, and I), HC area (B, F, and J), anterior commissure (AC) area (C, G, and K) or frequency of CC phenotypes (D, H, and L) in BTBR x C57 N2 mice. Genotyping for the 8 base pair deletion in *Draxin* in BTBR x C57 N2 mice compared with CC length (M), HC area (N), AC area (O), or frequency of CC phenotypes (P). Mice homozygous for the BTBR allele (C/C at SNP rs6397070 on chromosome (Chr) 4) or the *Draxin* deletion have significantly reduced CC length compared to heterozygous littermates. Data is represented as mean ± SEM. *** p < 0.001, **** p < 0.0001, ns = not significant, as determined by Kruskal-Wallis ANOVA tests. See related Figure S3.

### The Draxin deletion is associated with complete and partial CCD in BTBR x C57 N2 mice

To further demonstrate linkage of the eight-base pair deletion in *Draxin* to the observed CCD in BTBR and BTBR x C57 N2 mice, we performed Sanger sequencing of a SNP (rs6397070) on chromosome 4, 7.795 megabases downstream of the *Draxin* deletion as a marker for the BTBR allele in a subset of complete CCD, partial CCD and normal CC littermates (n ~ 10 of each phenotype, 31 total). Two additional SNPs were also sequenced at candidate loci on chromosome 9 (rs29890894) and chromosome 15 (rs31781085), which were previously judged to have a potential genetic influence on commissure size based on LOD score that did not reach statistical significance (Jones-Davis et al., 2013). We compared commissure length to the allele composition at the chromosome 4 locus (rs6397070), and found that the CC length was significantly reduced in mice homozygous for the BTBR allele (C/C) compared to heterozygous mice (C/T; Figure 5A and Table S1). In contrast, neither HC area (Figure 5B) or length (Figure S3A), or anterior commissure area (Figure 5C) demonstrated a significant difference between homozygous and heterozygous mice (Table S1), suggesting the chromosome 4 locus may determine variable CC formation in these mice. Of the 21 homozygous mice, 19 had CCD (n = 10 complete CCD, n = 9 partial CCD) and 2 had a normal CC. Of the 10 mice heterozygous at the rs6397070 allele, 9 had a normal CC and one mouse had partial CCD. No significant differences between genotypes were identified for chromosome 9 (Figure 5E-G, Figure S3B and Table S1) or chromosome 15 (Figure 5I–K, Figure S3C and Table S1) candidate loci for any commissure.

To further validate that the *Draxin* mutation is a likely cause of CCD in BTBR x C57 N2 mice, we genotyped for the eight-base pair *Draxin* deletion and examined the influence of the mutation on commissure size. CC length was significantly reduced in BTBR x C57 N2 mice homozygous for the *Draxin* deletion (Figure 5M and Table S1), consistent with our results of the candidate SNP on chromosome 4. All 16 mice from a cohort of 68 mice that displayed complete CCD were homozygous for the *Draxin* deletion (Figure 5P and Table S1). Of the 47 mice with normal CC length, 11 were homozygous for the *Draxin* deletion, suggesting incomplete penetrance of the mutation (Figure 5P and Table S1), which is also observed in *Draxin* knockout mice (Hossain et al., 2013; Islam et al., 2009). In line with the hypothesis that *Draxin* regulates the extent of IHF remodelling, which also influences HC formation, homozygosity for the *Draxin* deletion also significantly impacted HC length (Figure S3D and Table S1), but not HC area or anterior commissure area (Figure 5N, 50 and Table S1). Together, these results are consistent with a mutation in *Draxin* being the primary genetic aetiology underlying CCD and reduced HC length in the BTBR x C57 N2 mouse.

### Increased somal translocation of MZG and failure to remodel the IHF are associated with CCD in the BTBR strain

To investigate how DRAXIN regulates CC and HC formation in these mice, we first examined cell-type-specific *Draxin* expression in wildtype CD1 mice. *In situ* hybridization for *Draxin* with immunohistochemistry for glial markers and components of the IHF was performed prior to the onset of IHF remodelling. *Draxin* mRNA was highly expressed in GLAST-positive radial MZG progenitors in the telencephalic hinge from E12, and was further observed in GLAST-positive MZG migrating to the Laminin-positive IHF surface at E15 (Figure 6B and 6E). Immunohistochemistry on wildtype E15 horizontal sections revealed that DRAXIN was widely localized within the telencephalic midline on GLAST-positive radial MZG membranes including migrating cells and progenitors at all stages (Figure 6C, F, K), on innervating commissural axons at E15 (Figure 6H), and on the basement membrane of the IHF (Figure 6H’), and within the IHF (Figure 6H’). Thus, DRAXIN, which is known to be secreted (Islam et al., 2009), is expressed within MZG cells and associates with multiple cellular components of the interhemispheric midline, such that it could regulate the development of MZG, leptomeninges and axons, and the interactions between these cellular populations for IHF remodelling and corpus callosum formation.

**Figure 6:**
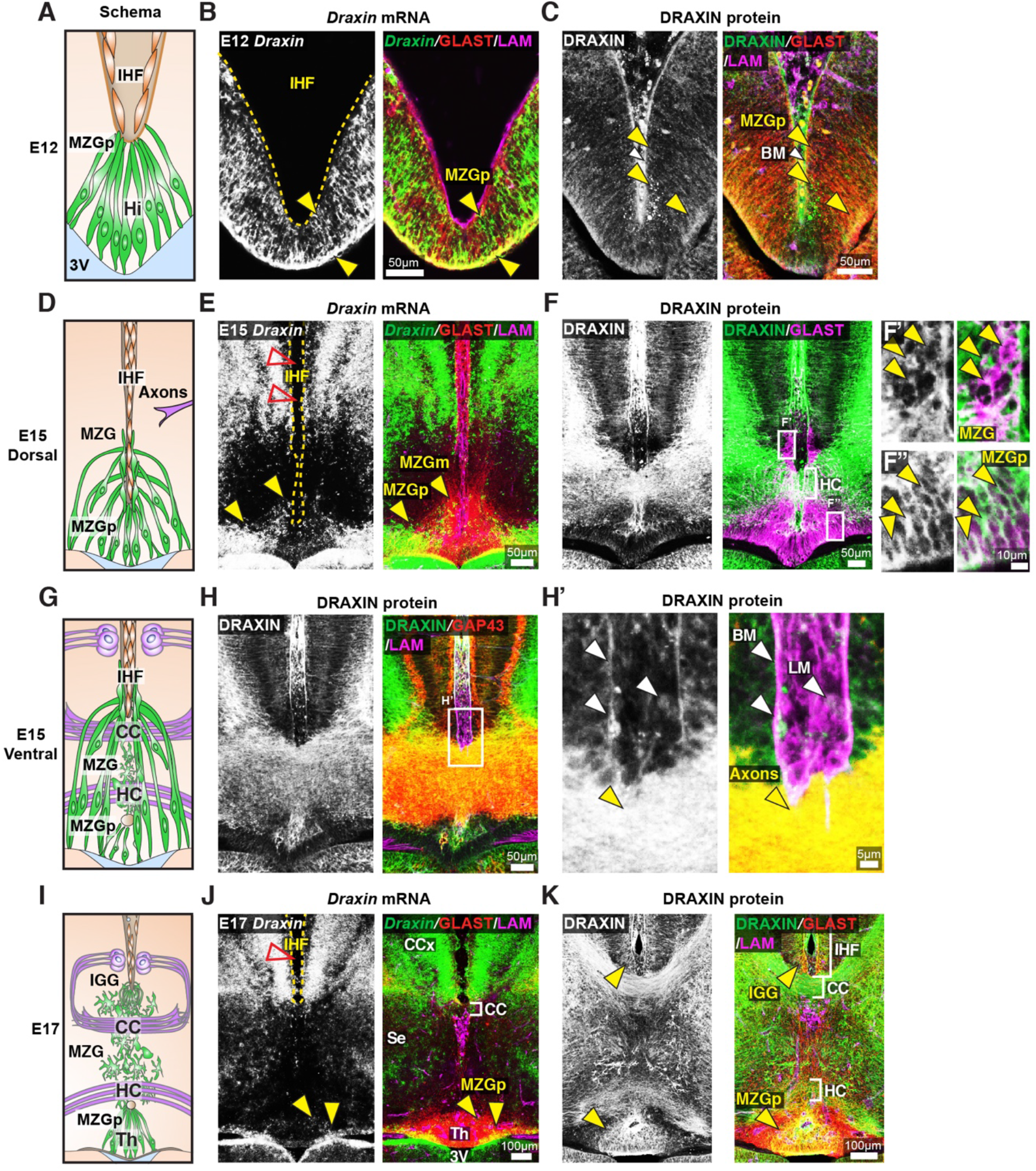
*Draxin* is expressed in MZG and their progenitors, and associates with MZG membranes, the pial surface of the IHF and leptomeninges. Schema of interhemispheric midline at E12 (A), E15 (D dorsal; G ventral) and E17 (I). *In situ* hybridization for *Draxin* mRNA (white or green), with immunohistochemistry for astroglial marker, GLAST (red), and leptomeninges and IHF marker, Laminin (LAM; magenta) in E12 (B), E15 (E) and E17 (J) wildtype CD1 mid-horizontal telencephalic midline tissue sections. Yellow arrowheads indicate *Draxin*-positive/GLAST-positive glia. Open red arrowheads indicate lack of *Draxin* mRNA within the IHF (yellow outlined). Immunohistochemistry for DRAXIN (white or green), GLAST (red or magenta), and LAM (magenta) in E12 (C), E15 (F) and E17 (K) wildtype CD1 mid-horizontal telencephalic midline tissue sections. (H) DRAXIN (white or green), axonal marker GAP43 (red) and LAM (magenta) in E15 ventral telencephalic midline tissue sections. Yellow arrowheads indicate regions of DRAXIN protein on GLAST-positive glial fibres (C, F and K) or DRAXIN protein on GAP43-positive axons (H’). White arrowheads indicate DRAXIN protein within the IHF and on the basement membrane of the IHF. BM = basement membrane, CCx = cingulate cortex, IGG = indusium griseum glia, LM = leptomeninges, MZG = midline zipper glia, MZGp = midline zipper glia progenitors, Se = septum, Th = telencephalic hinge, 3V = third ventricle.

To further investigate the function of DRAXIN during CC formation, we probed for cellular phenotypes that may explain the loss of IHF remodelling and CC formation in the BTBR parental strain. Immunohistochemistry for the leptomeningeal marker LAMININ (Gobius et al., 2016), axonal marker GAP43, and mature astroglial marker GFAP in E17 inbred BTBR and wildtype C57 mice revealed that BTBR inbred mice display complete retention of the IHF, which manifests as a significantly higher ratio of IHF length to total midline length in BTBR mice compared to wildtype C57 mice (Figure 7A-C and Table S1). GFAP-positive MZG remain in two columns of cells that do not intercalate across the IHF in BTBR inbred mice (Figure 7A), suggesting that a defect in the astroglial-IHF interaction results in the failure of midline crossing of CC and HC axons in BTBR mice.

**Figure 7:**
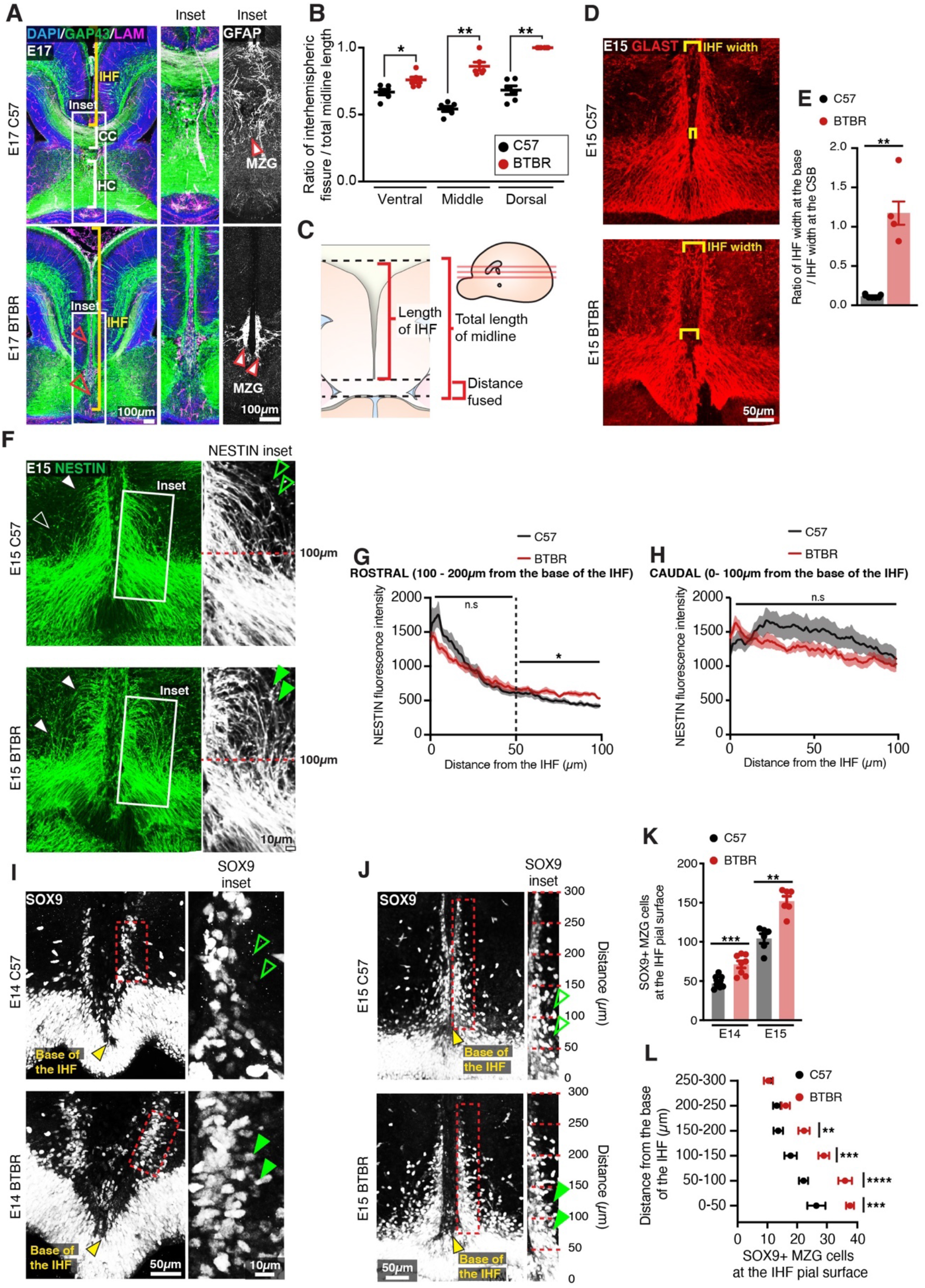
BTBR MZG undergo precocious somal translocation to the IHF and fail to intercalate for IHF remodelling. (A) Mid-horizontal sections of E17 wildtype C57 and BTBR mice immunolabelled with growing axon marker, GAP43 (green), astrocyte marker, GFAP (white; inset only), and leptomeninges marker, LAMININ (magenta), and counterstained with DAPI (blue). The CC and HC are indicated with white brackets in C57 mice, and their absence in BTBR mice is indicated with red arrowheads. The IHF is indicated with yellow brackets and white boxes indicate the region where insets of GFAP-positive MZG were taken (right). (C) The ratio of IHF length over total telencephalic midline length was measured (B) from representative ventral, middle (shown in A), and dorsal horizontal sections. Immunohistochemistry on E15 wildtype C57 and BTBR horizontal brain sections labelling GLAST-positive MZG (D) and NESTIN-positive radial glia (F) at the ventral midline. IHF width is indicated with yellow brackets in D and the ratio of IHF length close to the base of the IHF compared with at the corticoseptal boundary (CSB) is quantified in E. White arrowheads in F show radial MZG undergoing somal translocation to the IHF surface and green arrowheads in insets demonstrate an increase in radial MZG fibres lateral to the base of the IHF in BTBR mice; the fluorescence intensity of these NESTIN fibres is quantified in G (rostral to the base of the IHF) and H (caudal, close to the base of the IHF). Immunohistochemistry on E14 (I) and E15 (J) wildtype C57 and BTBR horizontal brain sections labelling SOX9-positive glial cell bodies. Green arrowheads indicate increased SOX9-positive cell bodies at the IHF surface in BTBR mice. The base of the IHF is indicated with yellow arrowheads. The number of MZG cell bodies at the pial surface (outlined in red) was quantified in K and L (binned). Data represent mean ± SEM. * p <0.05, ** p < 0.01, *** p < 0.001, **** p < 0.0001 as determined with either an unpaired t test (E14, K), Mann-Whitney test (E15, K) or 2-way ANOVA with Sidak’s multiple comparison test (L).

The endfeet of radial MZG progenitors normally form attachments to both the third ventricle and IHF within a region known as the telencephalic hinge. They then proliferate and undergo somal translocation to the IHF between E12 and E15 to initiate IHF remodelling at E15 in mice (Gobius et al., 2016). Immunohistochemistry for radial MZG using NESTIN and GLAST prior to IHF remodelling revealed that while radial MZG are present between the IHF pial surface and the third ventricle in BTBR mice (Figure 7D and F), radial NESTIN-positive MZG fibres were disorganised, and significantly more abundant lateral to the IHF between 100 – 200 μm from the base of the IHF in BTBR mice (Figure 7F, 7G and Table S1). Moreover, altered radial MZG distribution in BTBR mice was associated with abnormal widening of the base of the IHF (Figure 7D-E and Table S1); a region that normally undergoes selective compression prior to IHF remodelling as increasing numbers of MZG translocate to it (Gobius et al., 2016). Immunohistochemistry for radial glia and astrocyte marker, SOX9 (Sun et al., 2017), revealed significantly increased SOX9-positive MZG cell bodies at the pial surface of the IHF in the BTBR inbred strain at E14 and E15 compared to wildtype C57 mice (Figure 7I-K and Table S1). These MZG clustered along the IHF surface within 200 μm of the base of the IHF at E15 (Figure 7L and Table S1), but did not result in narrowing of the adjacent leptomeninges, indicating that earlier defects in the generation of the MZG population could underlie the persistence of the IHF in BTBR mice.

### Increased proliferation and cell cycle exit of MZG progenitors is associated with increased somal translocation of MZG and may underlie CC agenesis in the parental BTBR strain

We have previously demonstrated increased proliferation of radial glia within the E14 cingulate cortex of the BTBR inbred mouse strain (Fariday et al., 2014). To determine whether a similar effect may underlie increased somal translocation of MZG in BTBR mice, we performed birth-dating of MZG progenitors by EdU injections and co-labelling with cell cycle marker, KI67. This analysis revealed an increased division of MZG progenitors in BTBR mice at E13 and E14 leading to significantly increased labelling of EdU-positive MZG at E14 and E15, respectively (Figure 8A-D, 8F and Table S1). Between E14 and E15, more dividing MZG progenitors exited the cell cycle (EdU-positive/KI67-negative; Figure 8H and Table S1) in BTBR mice compared with C57 controls. These differences were unlikely to be due to mismatched developmental stage of the embryos since the length of the telencephalic midline was comparable between BTBR and wildtype C57 brains at the age of collection (Figure 8E and Table S1). We find increased proliferation and cell cycle exit of MZG progenitors in BTBR mice leads to an over-abundance of NESTIN-positive MZG fibres and increased somal translocation of MZG to the IHF surface prior to IHF remodelling. Since we previously found that precocious generation of MZG is associated with disrupted IHF remodelling in mice with altered FGF8 signalling (Gobius et al., 2016), our results here suggest that disrupted IHF remodelling in BTBR mice is likely due to precocious generation of MZG.

**Figure 8:**
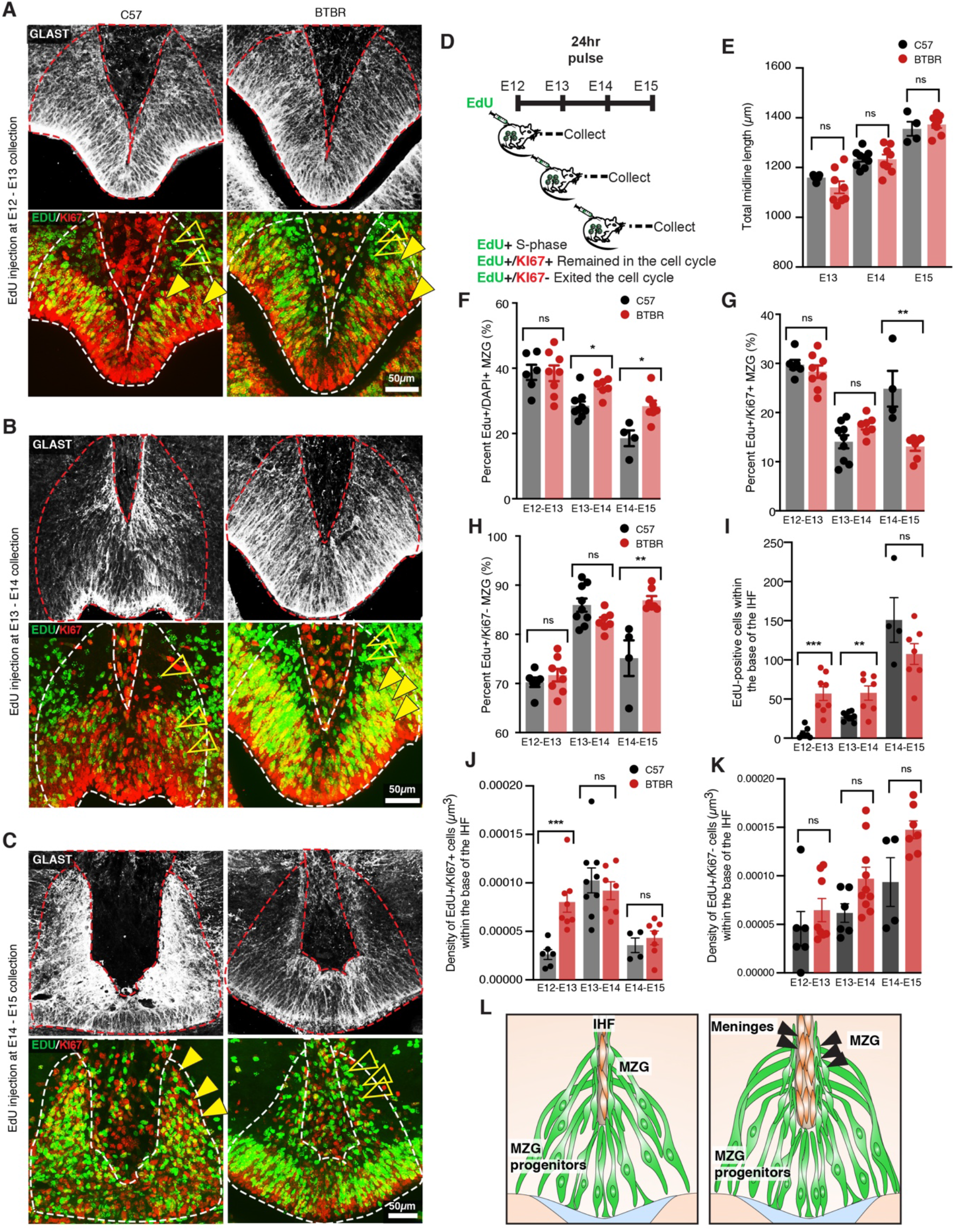
Elevated MZG progenitor and leptomeningeal proliferation in BTBR mice. Wildtype C57 and BTBR pregnant mice were injected with EdU every 24 hours from E12 and collected 24 hours later (D). Immunohistochemistry for GLAST (white), EdU (green) and cell cycle marker, KI67 (red) on E13 (A), E14 (B) and E15 (C) wildtype C57 and BTBR horizontal brain sections of the telencephalic hinge and IHF base. (E) To determine whether the litters were age-matched, the total length of the midline was compared between groups. The percentage of EdU-positive/DAPI-positive, EdU-positive/KI67-positive, and EdU-positive/KI67-negative MZG from the telencephalic hinge (white dotted outline) is quantified in F, G and H respectively. EdU-positive cells within the base of the IHF is quantified in I. EdU-positive/KI67-positive cells or EdU-positive/KI67-negative cells within the IHF were normalised to the total volume of the IHF as quantified in J and K, respectively. Data represent mean ± SEM, * p < 0.05, ** p < 0.01, *** p < 0.001, ns = not significant, as determined with Mann-Whitney tests. (L) Schema of BTBR phenotype at E15 compared with wildtype C57: BTBR mice display increased proliferation of MZG progenitors and precocious migration to the IHF surface as well as proliferation of the leptomeninges and expansion of the IHF, which may underlie failed IHF remodelling in these mice. See related Figure S3.

We further measured the proliferative behaviour of cells within the base of the IHF that is surrounded by the telencephalic hinge, and found a significant increase in EdU-positive cells at E13 and E14, which were dividing at E12 and E13, respectively, in BTBR mice (Figure 8A-B, 8I and Table S1). The density of cells that remained in the cell cycle after dividing was significantly increased between E12 and E13 in BTBR mice (EdU-positive/KI67-positive; Figure 8A, 8K and Table S1). There were no differences in the density of cells that exited the cell cycle within the BTBR IHF compared to the wildtype C57 IHF (EdU-positive/KI67-negative; Figure 8A-C, 8K and Table S1). Cells within the IHF comprise CXCL12- and LAMININ-positive leptomeningeal cells, which are eliminated from the septum during IHF remodelling (Gobius et al., 2016). These findings suggest that failed IHF remodelling in BTBR mice is correlated with a transient period of increased proliferation of both leptomeningeal cells within the IHF and MZG progenitors, and subsequent over-generation and migration of MZG to an enlarged IHF in BTBR mice.

## Discussion

Altogether, this work provides a comprehensive view of the genetic and cellular aetiology of CCD in a mouse model of variable forebrain commissure dysgenesis. It reveals that an eight-base pair deletion in *Draxin* affecting protein structure and expression is the main genetic driver of malformations of the CC and HC in BTBR and BTBR x C57 N2 mice. Moreover, abnormal proliferation of MZG progenitors and leptomeningeal cells within the IHF, and subsequent surplus migration of MZG was associated with aberrant remodelling of the IHF; the extent of which was further correlated with the severity of commissure dysgenesis in outcrossed BTBR strains. Moreover, evidence of incomplete septal fusion and IHF remodelling in a cohort of human individuals with partial CCD, suggests a common structural aetiology for a wide range of human CCD phenotypes.

The developmental basis of varied severity and expressivity of CCD in humans has to date been unclear. For example, humans with pathogenic variants in the DRAXIN receptor, *DCC*, display a range of CC phenotypes and associated HC malformations even across family members carrying the same pathogenic variant (Jamuar et al., 2017; Marsh et al., 2017; Marsh et al., 2018). *Dcc* mutant mice, which carry loss of function mutations, do not recapitulate the phenotypic variability observed in humans (Fazeli et al., 1997; Finger et al., 2002; Fothergill et al., 2013). Here, we demonstrate that the severity of commissural dysgenesis can be explained by the extent to which the IHF has failed to be remodelled. This in turn, determines the volume of substrate formed to allow axons to cross the interhemispheric midline. Non-axonal intrinsic mechanisms therefore determine the severity of the phenotype, but axon guidance defects may also contribute. Exemplified by the phenotype of BTBR x C57 N2 mice with partial CCD, it appears that even when carrying the same genetic disruption, callosal axons may still cross the midline where there is a permissive substrate available. Genetic perturbation that affects axon guidance mechanisms is therefore likely to be malleable and less disruptive for CC formation. In contrast, any disruption to IHF remodelling is likely irrevocable, and our findings indicate it is the main determinant of a spectrum of CCD. Whether other genetic or environmental modifiers exist to determine the success of IHF remodelling remains an important question for future research. While pathogenic variants in *DRAXIN* have not yet been reported in human individuals with CCD, a similar effect may underlie the spectrum of phenotypes observed in humans with *DCC* mutations, since *Dcc* and *Draxin* have been demonstrated to interact to determine the severity of CCD in mice (Ahmed et al., 2011).

Failure for a midline substrate to form has been previously associated with incomplete penetrance of CCD and HC dysgenesis in BALB/cWah1 and 129P1/ReJ inbred mouse strains (Bohlen et al., 2012; Wahlsten et al., 2006). Importantly, in BTBR x C57 N2 mice, we found HC length was significantly reduced in mice with CCD compared to full CC littermates, was reduced in complete CCD mice compared to partial CCD mice, and was significantly reduced in mice homozygous for the *Draxin* deletion. Together, these results are consistent with a similar primary developmental defect underlying HC dysgenesis and CCD. To our knowledge, this is also the first description of incomplete IHF remodelling in humans with partial CCD. Thus, failure to remodel the IHF underlies not just complete CCD, but a spectrum of CCD and HC dysgenesis. IHF remodelling thus appears to be a continuous process that can be variably disrupted; the cellular and molecular pathways that determine its success remain to be investigated.

DRAXIN has been classically described as an axon guidance ligand, which acts as a chemorepulsive cue to axons derived from cortical explants (Islam et al., 2009). *Draxin* knockout mice, however, display variable penetrance of CCD (Ahmed et al., 2011; Islam et al., 2009; Hossain et al., 2013). It is therefore unsurprising that homozygous inheritance of the *Draxin* deletion in BTBR x C57 N2 mice yielded approximately 60% of mice with CCD and 40% of mice with a normal CC, since the same penetrance has been observed in *Draxin* knockout mice maintained on a C57 genetic background (Hossain et al., 2013). This is consistent with the idea that the *Draxin* deletion in BTBR mice is a loss of function mutation that drives CC abnormalities at incomplete penetrance. A similar locus on chromosome 4 has previously been implicated in CC size of an intercross between NZB/BINJ and C57Bl/6By mice (Roy, Perez-Diaz and Roubertoux, 1998), suggesting that mutations influencing *Draxin* function may underlie CCD in at least two inbred mouse strains.

Here we show a distinct, earlier role for DRAXIN during corpus callosum formation in the astroglial-dependent formation of an interhemispheric substrate. *Draxin* expression within the MZG cell lineage likely influences MZG cellular proliferation and migration to the interhemispheric midline for remodelling, since these cellular processes were disrupted in BTBR mice. We previously demonstrated that dysfunction of these processes within MZG are causally associated with failed IHF remodelling in mice with altered expression of astrogliogenesis factors, FGF8, NFIA, and NFIB (Gobius et al., 2016). Whether DRAXIN acts downstream of FGF8-NFI signalling to regulate MZG development and IHF remodelling could be investigated in a future study. Further, we found DRAXIN protein within the IHF, which was negative for *Draxin* mRNA, suggesting secreted DRAXIN may cross from the neural to a non-neural compartment. DRAXIN is known to regulate the migration of cranial neural crest cells at earlier stages of development (Hutchins and Bronner, 2018), and is also involved in basement membrane remodelling during cranial neural crest epithelial to mesenchyme transition in chicks (Hutchins and Bronner, 2019). Considering these roles, DRAXIN may directly regulate the leptomeninges (which are thought to arise from the neural crest; Batarfi et al., 2017), or IHF remodelling, possibly by promoting repulsive migration of leptomeninges from the midline or remodelling the basement membrane of the IHF. Indeed, we observed an abnormally large IHF in BTBR mice at the time of CC development that was directly associated with the increased proliferation of cells within the base of the IHF. DRAXIN is known to interact with the canonical WNT receptor LRP6, and antagonize canonical WNT signalling (Miyake et al., 2009). Moreover, canonical WNT signalling within cortical radial glia controls cell proliferation and astrogliogenesis (Gan et al., 2014). Thus, if LRP6 is also expressed by MZG or leptomeningeal cells, it may be the molecular link between DRAXIN and the regulation of cell proliferation and elevated generation of MZG and leptomeninges, which we observed in BTBR mice. Together, precocious MZG migration and increased abundance of leptomeninges within the IHF could prevent MZG intercalation and IHF remodelling due to the increased distance between ipsilateral and contralateral MZG processes. Thus, DRAXIN, which is thought to influence CC formation via axon intrinsic mechanisms (Ahmed et al., 2011; Edwards et al, 2014; Islam et al., 2009; Morcom et al., 2016), may rather influence several cellular processes in the telencephalic midline required for IHF remodelling and commissure formation. Importantly, the cumulative success of multiple processes under the control of DRAXIN may be crucial for normal CC development, while dysfunction of one or more of these processes may instead determine the severity of the callosal phenotype. Conditional strategies that impact DRAXIN function in axons versus MZG versus leptomeninges are necessary to dissect these possibilities.

## Acknowledgements

We thank Aiman Al Najjar, Nicole Atcheson and Dr. Nyoman Kurniawan for assistance in conducting MRI at the Centre for Advanced Imaging, The University of Queensland. We thank Rumelo Amor, Arnaud Guardin and Andrew Thompson for assistance with microscopy, which was performed in the Queensland Brain Institute’s Advanced Microscopy Facility. We thank the staff of the University of Queensland Biological Resources for care and breeding of animals. This work was supported by Australian NHMRC grants GNT1048849 and GNT1126153 to L.J.R and USA National Institutes of Health grant 5R01NS058721 to E.H.S and L.J.R. R.S received an Australian Research Council DECRA fellowship (DE160101394). L.M and J.W.C.L were supported by a Research training program scholarship (Australian Postgraduate Award). T.J.E and K.S.C were supported by a University of Queensland Research Scholarship. L.M, T.J.E, and J.W.C.L also received Queensland Brain Institute Top-Up scholarships. L.J.R. was supported by an NHMRC Principal Research Fellowship (GNT1120615).

We thank the families and members of the Australian Disorders of the Corpus Callosum (AusDoCC) for their support and time in being involved in this research. We thank the International Research Consortium for the Corpus Callosum and Cerebral Connectivity (IRC5, https://www.irc5.org) researchers for discussions and input.

## Notes

### Competing Interest Statement

The authors have declared no competing interest.

## References

Ahmed, G., Tessier-Lavigne, M., Tanaka, H., Shinmyo, Y., Ohta, K., Islam, S.M., Hossain, M., Naser, I.B., Riyadh, M.A., Su, Y., et al. (2011). Draxin inhibits axonal outgrowth through the netrin receptor DCC. The Journal of Neuroscience 31, 14018–14023.

Batarfi, M., Valasek, P., Krejci, E., Huang, R., and Patel, K. (2017). The development and origins of vertebrate meninges. Biological Communications 62, 73–81.

Bohlen, M.O., Bailoo, J.D., Jordan, R.L., and Wahlsten, D. (2012). Hippocampal commissure defects in crosses of four inbred mouse strains with absent corpus callosum. Genes, Brain and Behavior 11, 757–766.

Brown, W.S., and Paul, L.K. (2019). The neuropsychological syndrome of agenesis of the corpus callosum. Journal of the International Neuropsychological Society, 1–7.

Bunt, J., de Haas, T.G., Hasselt, N.E., Zwijnenburg, D.A., Koster, J., Versteeg, R., and Kool, M. (2010). Regulation of cell cycle genes and induction of senescence by overexpression of OTX2 in medulloblastoma cell lines. Molecular Cancer Research 8, 1344.

Bunt, J., Osinski, J.M., Lim, J.W.C., Vidovic, D., Ye, Y., Zalucki, O., O’Connor, T.R., Harris, L., Gronostajski, R.M., Richards, L.J., et al. (2017). Combined allelic dosage of Nfia and Nfib regulates cortical development. Brain and Neuroscience Advances 1, 2398212817739433.

Chen, K.-S., Bridges, C.R., Lynton, Z., Lim, J.W.C., Stringer, B.W., Rajagopal, R., Wong, K.-T., Ganesan, D., Ariffin, H., Day, B.W., et al. (2020). Transcription factors NFIA and NFIB induce cellular differentiation in high-grade astrocytoma. Journal of Neuro-Oncology 146, 41–53.

Donahoo, A.-L.S., and Richards, L.J. (2009). Understanding the mechanisms of callosal development through the use of transgenic mouse models. Paper presented at: Seminars in Pediatric Neurology (Elsevier).

Edwards, T.J., Sherr, E.H., Barkovich, A.J., and Richards, L.J. (2014). Clinical, genetic and imaging findings identify new causes for corpus callosum development syndromes. Brain 137, 1579–1613.

Edwards, T. J., Fenlon, L. R., Dean, R. J., Bunt, J., Sherr, E. H., & Richards, L. J. (2020). Altered structural connectivity networks in a mouse model of complete and partial dysgenesis of the corpus callosum. NeuroImage, 217, 116868.

Fazeli, A., Dickinson, S.L., Hermiston, M.L., Tighe, R.V., Steen, R.G., Small, C.G., Stoeckli, E.T., Keino-Masu, K., Masu, M., and Rayburn, H. (1997). Phenotype of mice lacking functional deleted in colorectal cancer (Dec) gene. Nature 386, 796.

Finger, J.H., Bronson, R.T., Harris, B., Johnson, K., Przyborski, S.A., and Ackerman, S.L. (2002). The netrin 1 receptors Unc5h3 and Dcc are necessary at multiple choice points for the guidance of corticospinal tract axons. Journal of Neuroscience 22, 10346–10356.

Fothergill, T., Donahoo, A.-L.S., Douglass, A., Zalucki, O., Yuan, J., Shu, T., Goodhill, G.J., and Richards, L.J. (2013). Netrin-DCC signaling regulates corpus callosum formation through attraction of pioneering axons and by modulating Slit2-mediated repulsion. Cerebral Cortex 24, 1138–1151.

Glass, H.C., Shaw, G.M., Ma, C., and Sherr, E.H. (2008). Agenesis of the corpus callosum in California 1983–2003: A population-based study. American Journal of Medical Genetics Part A 146, 2495–2500.

Gobius, I., Morcom, L., Suárez, R., Bunt, J., Bukshpun, P., Reardon, W., Dobyns, William B., Rubenstein, John L.R., Barkovich, A.J., Sherr, Elliott H., et al. (2016). Astroglial-mediated remodeling of the interhemispheric midline is required for the formation of the corpus callosum. Cell Reports 17, 735–747.

Hearne, L. J., Dean, R. J., Robinson, G. A., Richards, L. J., Mattingley, J. B., and Cocchi, L. (2019). Increased cognitive complexity reveals abnormal brain network activity in individuals with corpus callosum dysgenesis. NeuroImage: Clinical 21, 101595.

Hetts, S.W., Sherr, E.H., Chao, S., Gobuty, S., and Barkovich, A.J. (2006). Anomalies of the corpus callosum: an MR analysis of the phenotypic spectrum of associated malformations. American Journal of Roentgenology 187, 1343–1348.

Hutchins, E.J., and Bronner, M.E. (2018). Draxin acts as a molecular rheostat of canonical Wnt signaling to control cranial neural crest EMT. Journal of Cell Biology 217, 3683–3697.

Hutchins, E.J., and Bronner, M.E. (2019). Draxin alters laminin organization during basement membrane remodeling to control cranial neural crest EMT. Developmental Biology 446, 151–158.

Islam, S.M., Shinmyo, Y., Okafuji, T., Su, Y., Naser, I.B., Ahmed, G., Zhang, S., Chen, S., Ohta, K., Kiyonari, H., et al. (2009). Draxin, a repulsive guidance protein for spinal cord and forebrain commissures. Science (New York, NY) 323, 388–393.

Jamuar, S.S., Schmitz-Abe, K., D’Gama, A.M., Drottar, M., Chan, W.-M., Peeva, M., Servattalab, S., Lam, A.-T.N., Delgado, M.R., and Clegg, N.J. (2017). Biallelic mutations in human DCC cause developmental split-brain syndrome. Nature Genetics 49, 606.

Lein, E.S., Hawrylycz, M.J., Ao, N., Ayres, M., Bensinger, A., Bernard, A., Boe, A.F., Boguski, M.S., Brockway, K.S., and Byrnes, E.J. (2007). Genome-wide atlas of gene expression in the adult mouse brain. Nature 445, 168.

Marsh, A.P., Edwards, T.J., Galea, C., Cooper, H.M., Engle, E.C., Jamuar, S.S., Méneret, A., Moutard, M.L., Nava, C., and Rastetter, A. (2018). DCC mutation update: Congenital mirror movements, isolated agenesis of the corpus callosum, and developmental split brain syndrome. Human Mutation 39, 23–39.

Marsh, A.P., Heron, D., Edwards, T.J., Quartier, A., Galea, C., Nava, C., Rastetter, A., Moutard, M.-L., Anderson, V., and Bitoun, P. (2017). Mutations in DCC cause isolated agenesis of the corpus callosum with incomplete penetrance. Nature Genetics 49, 511.

Moldrich, R.X., Gobius, I., Pollak, T., Zhang, J., Ren, T., Brown, L., Mori, S., De Juan Romero, C., Britanova, O., Tarabykin, V., and Richards, L.J. (2010). Molecular regulation of the developing commissural plate. Journal of Comparative Neurology 518, 3645–3661.

Morcom, L.R., Edwards, T.J., and Richards, L.J. (2015). Cortical architecture, midline guidance, and tractography of 3D white matter tracts. In Axons and brain architecture, K.S. Rockland, ed. (Cambridge, United Kingdom: Academic Press), pp. 289–313.

Paul, L.K., Brown, W.S., Adolphs, R., Tyszka, J.M., Richards, L.J., Mukherjee, P., and Sherr, E.H. (2007). Agenesis of the corpus callosum: genetic, developmental and functional aspects of connectivity. Nature reviews Neuroscience 8, 287–299.

Probst, M. (1901). The structure of complete dissolving corpus callosum of the cerebrum and also the microgyry and heterotropy of the grey substance. Arch Psychiatr Nervenkr 34.

Rakic, P., and Yakovlev, P.I. (1968). Development of the corpus callosum and cavum septi in man. The Journal of Comparative Neurology 132, 45–72.

Schanze, I., Bunt, J., Lim, J.W.C., Schanze, D., Dean, R.J., Alders, M., Blanchet, P., Attié-Bitach, T., Berland, S., Boogert, S., et al. (2018). NFIB Haploinsufficiency Is Associated with Intellectual Disability and Macrocephaly. The American Journal of Human Genetics 103, 752–768.

Silver, J., Edwards, M.A., and Levitt, P. (1993). Immunocytochemical demonstration of early appearing astroglial structures that form boundaries and pathways along axon tracts in the fetal brain. The Journal of Comparative Neurology 328, 415–436.

Suárez, R., Fenlon, L.R., Marek, R., Avitan, L., Sah, P., Goodhill, G.J., and Richards, L.J. (2014). Balanced interhemispheric cortical activity is required for correct targeting of the corpus callosum. Neuron 82, 1289–1298.

Ullmann, J. F. P., Watson, C., Janke, A. L., Kurniawan, N. D., and Reutens, D. C. (2013). A segmentation protocol and MRI atlas of the C57BL/6J mouse neocortex. NeuroImage, 78, 196–203.

Wahlsten, D., Bishop, K., and Ozaki, H. (2006). Recombinant inbreeding in mice reveals thresholds in embryonic corpus callosum development. Genes, Brain and Behavior 5, 170–188.

Wahlsten, D., Metten, P., and Crabbe, J.C. (2003). Survey of 21 inbred mouse strains in two laboratories reveals that BTBR T/+ tf/tf has severely reduced hippocampal commissure and absent corpus callosum. Brain Research 971, 47–54.

Yushkevich, P. A., Piven, J., Hazlett, H. C., Smith, R. G., Ho, S., Gee, J. C., and Gerig, G. (2006). User-guided 3D active contour segmentation of anatomical structures: significantly improved efficiency and reliability. Neuroimage, 31, 1116–1128.

