## Supplementary figures and table for "DRAXIN regulates interhemispheric fissure remodelling to influence the extent of corpus callosum formation"

**Supplementary figures and titles:**

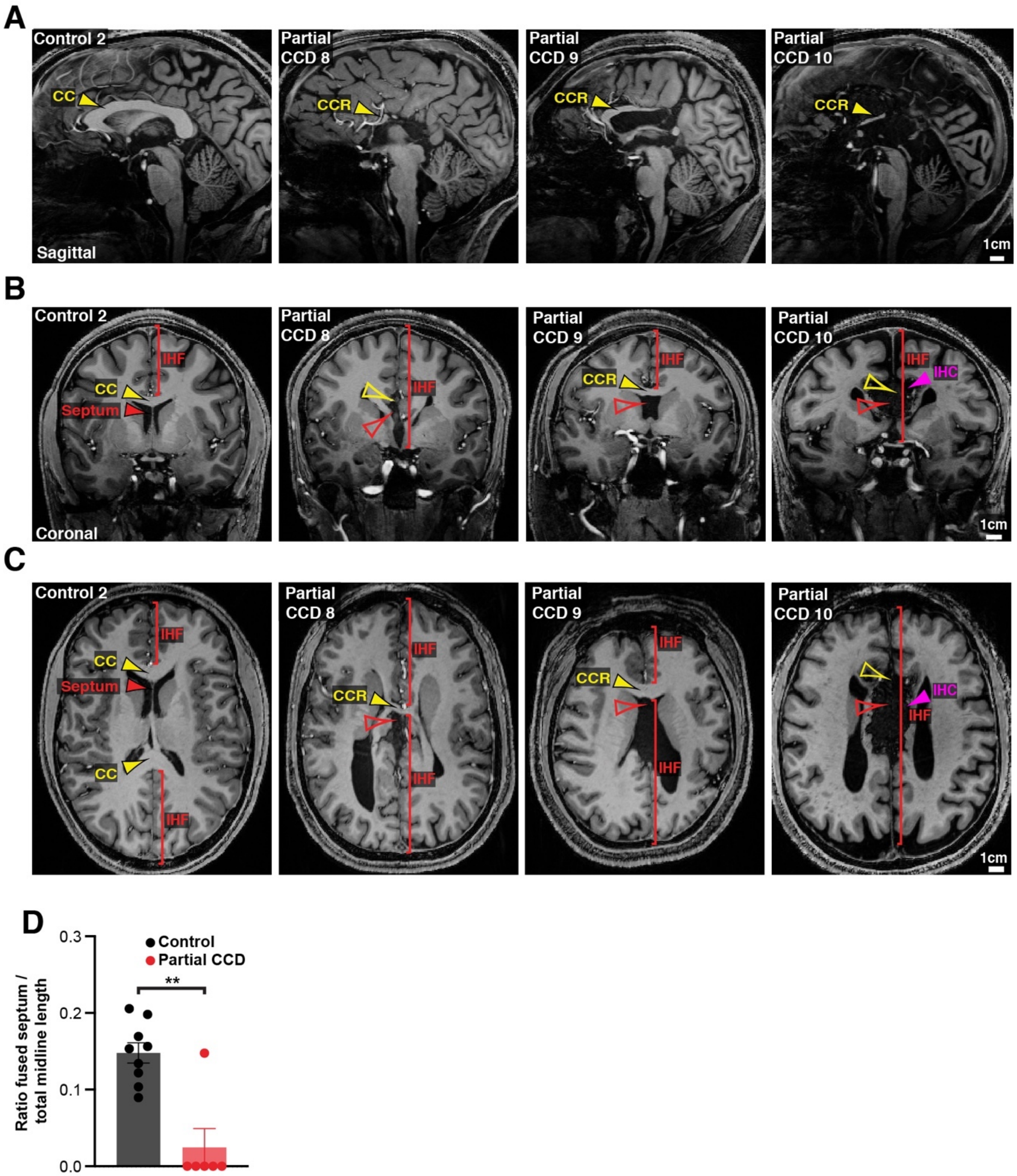


**Figure S1: Structural MRI study of IHF and septal defects associated with partial CCD in humans**

Sagittal (A), coronal (B) and axial (C) slices from T1-weighted structural scans from remaining partial CCD individuals (4/10) and another neurotypical (control) individual

(1/9). The CC or the CC remnant (CCR) is indicated with yellow arrowheads, presence of an interhemispheric cyst (IHC) is indicated with magenta arrowheads, the IHF extent is indicated with red brackets and the septum, septal leaves or absence of the septal substrate is indicated with red arrowheads. (D) The ratio of fused septum was measured from axial images in C (and Figure 3C) and normalised to the total midline length. Data is represented as mean ± SEM. ** $p < 0.01$, as determined with an unpaired t test. See related Figure 3 for further subjects and quantification.

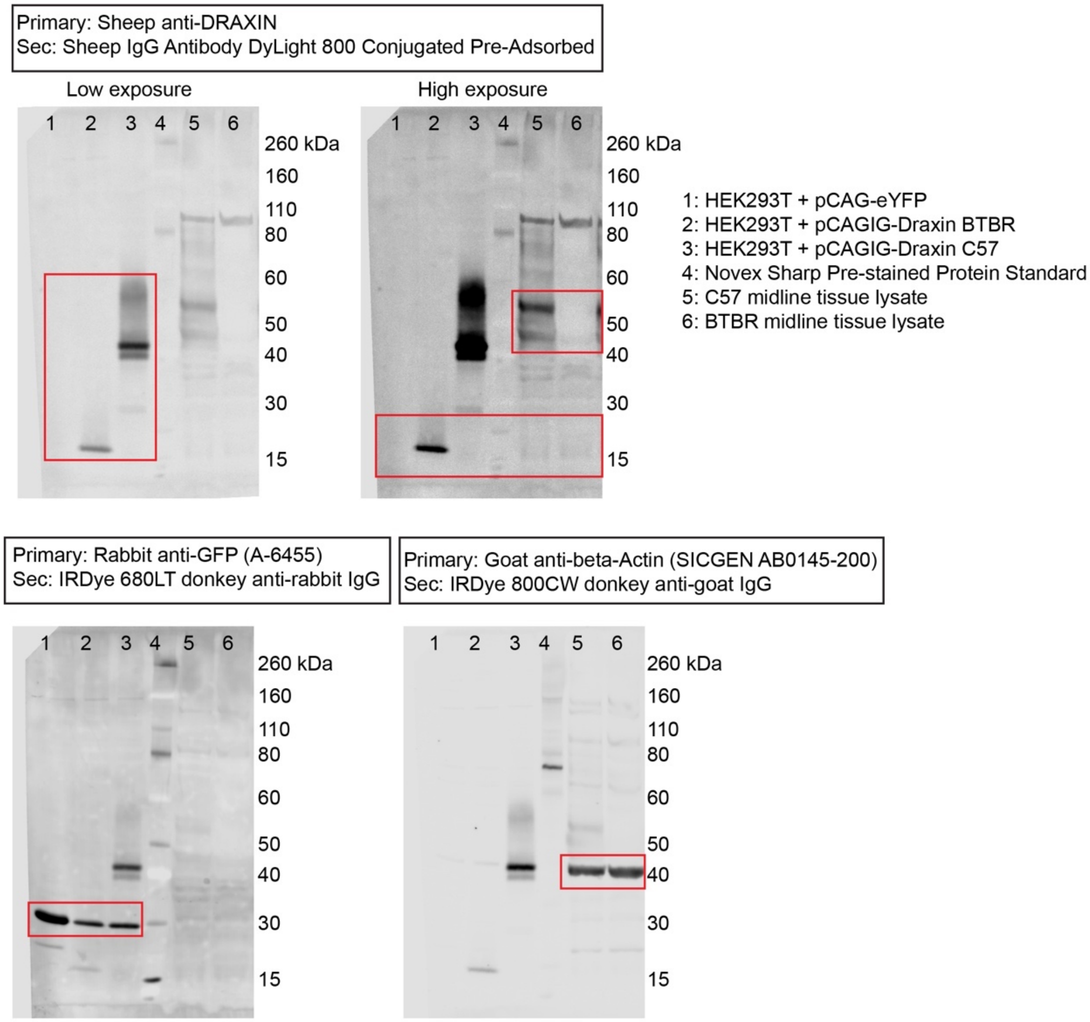


**Figure S2: Western blots for DRAXIN in HEK293T cell lysates expressing the coding sequence of BTBR or C57 *Draxin,* and in E15 midline tissue from C57 and BTBR mice**

Cell lysates derived from HEK293T cells expressing pCag-eYFP, or pCag-iresGFP with either BTBR or C57 *Draxin* coding sequences were incubated with anti-DRAXIN and anti-GFP antibodies. Specific bands at ~18kD and ~40-45kD demonstrate that BTBR *Draxin* produces a protein of reduced molecular weight, indicating truncation. Midline tissue lysates from E15 C57 and BTBR mice incubated with anti-DRAXIN and anti-β-ACTIN antibodies reveal specific bands at ~40-60kD and ~42kD that indicate DRAXIN expression is severely reduced in BTBR mice. Sec = secondary.

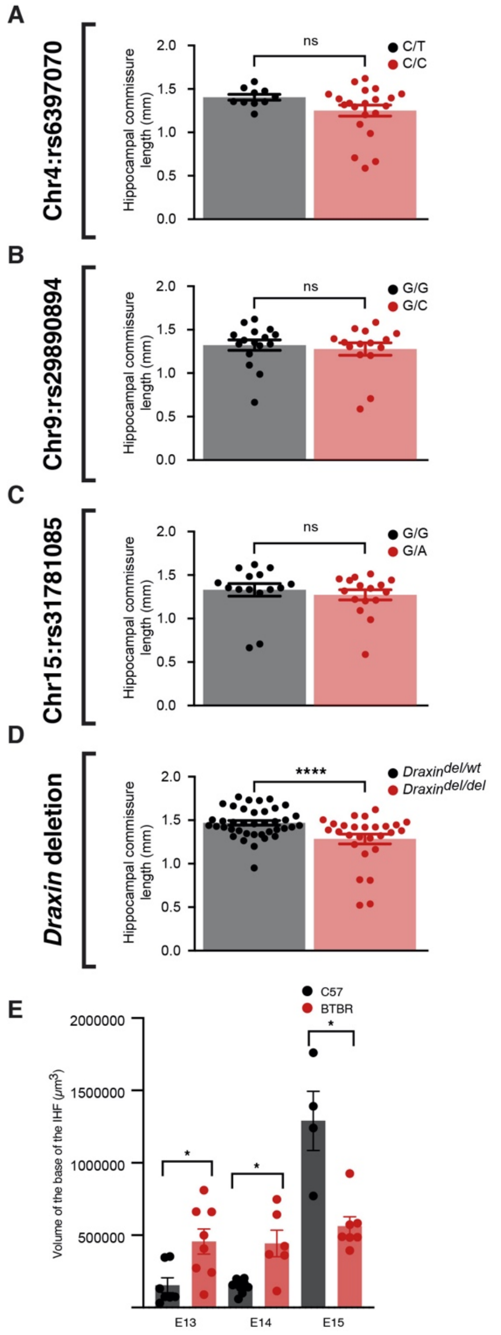

**Figure S3: Group-wise comparison of marker SNPs for candidate genetic loci and a *Draxin* mutation to explain HC length in BTBR x C57 N2 mice and volume of the IHF base in BTBR mice**

Group-wise comparison for SNPs at candidate genetic loci (A-C) or genotyping for the 8 base pair *Draxin* deletion (D) compared with HC length in BTBR x C57 N2 mice. (E)

Volume of the IHF base analysed in C57 and BTBR mice that was used to quantify EdU-positive cells in Figure 8I-K. * p < 0.05, ** p < 0.01, ** p < 0.001, **** p < 0.0001, ns = not significant, as determined by Kruskal-Wallis ANOVA tests (A-C) or Mann-Whitney tests (D-E). See related Figures 5 and 8.

### Supplementary Table 1: Statistics

| Figure | Data compared | Number of samples | Statistical test | p value |
|---|---|---|---|---|
| 1C | HC length cCCD vs pCCD | n = 27 cCCD, n = 15 pCCD | Welch's t test | 0.0126 |
| 1C | HC length cCCD vs full CC | n = 27 cCCD, n = 66 full CC | Welch's t test | < 0.0001 |
| 1C | HC length pCCD vs full CC | n = 15 pCCD, n = 66 full CC | Welch's t test | 0.0021 |
| 3D | Ratio dorso-ventral IHF length pCCD humans | n = 9 control, n = 10 pCCD | Unpaired t test | 0.0044 |
| 3E | Ratio dorso-ventral CC width pCCD humans | n = 9 control, n = 10 pCCD | Mann-Whitney test | 0.0322 |
| 3G | Ratio anterior-posterior IHF length anterior segment pCCD humans | n = 9 control, n = 8 pCCD | Unpaired t test | 0.0023 |
| 3G | Ratio anterior-posterior IHF length posterior segment pCCD humans | n = 9 control, n = 8 pCCD | Unpaired t test | <0.0001 |
| 3H | Ratio anterior-posterior CC length anterior segment pCCD humans | n = 9 control, n = 10 pCCD | Unpaired t test | 0.0029 |
| 3H | Ratio anterior-posterior CC length posterior segment pCCD humans | n = 9 control, n = 10 pCCD | Mann-Whitney test | < 0.0001 |
| 5A | CC length chr4:rs6397070 SNP | n = 10 heterozygous, n = 21 homozygous | Kruskal-Wallis ANOVA with Dunn's multiple comparison | 0.0006 |

| | | | | |
|---|---|---|---|---|
| 5B | HC area chr4:rs6397070 SNP | n = 10 heterozygous, n = 21 homozygous | Kruskal-Wallis ANOVA with Dunn's multiple comparison test | > 0.9999 |
| 5C | AC area chr4:rs6397070 SNP | n = 10 heterozygous, n = 21 homozygous | Kruskal-Wallis ANOVA with Dunn's multiple comparison test | > 0.9999 |
| 5E | CC length chr9:rs29890894 SNP | n = 15 heterozygous, n = 16 homozygous | Kruskal-Wallis ANOVA with Dunn's multiple comparison | > 0.9999 |
| 5F | HC area chr9:rs29890894 SNP | n = 15 heterozygous, n = 16 homozygous | Kruskal-Wallis ANOVA with Dunn's multiple comparison test | > 0.9999 |
| 5G | AC area chr9:rs29890894 SNP | n = 15 heterozygous, n = 16 homozygous | Kruskal-Wallis ANOVA with Dunn's multiple comparison test | > 0.9999 |
| 5I | CC length chr15:rs31781085 SNP | n = 16 heterozygous, n = 15 homozygous | Kruskal-Wallis ANOVA with Dunn's multiple comparison | > 0.9999 |
| 5J | HC area chr15:rs31781085 SNP | n = 16 heterozygous, n = 15 homozygous | Kruskal-Wallis ANOVA with Dunn's multiple comparison test | > 0.9999 |
| 5K | AC area chr15:rs31781085 SNP | n = 16 heterozygous, n = 15 homozygous | Kruskal-Wallis ANOVA with Dunn's multiple comparison test | > 0.9999 |
| 5M | CC length *Draxin* mutation | n = 39 heterozygous, n = 29 homozygous | Kruskal-Wallis ANOVA with Dunn's multiple comparison test | 0.0002 |

| | | | | |
|---|---|---|---|---|
| 5N | HC area *Draxin* mutation | n = 39 heterozygous, n = 29 homozygous | Kruskal-Wallis ANOVA with Dunn's multiple comparison test | > 0.9999 |
| 5O | AC area *Draxin* mutation | n = 39 heterozygous, n = 29 homozygous | Kruskal-Wallis ANOVA with Dunn's multiple comparison test | > 0.9999 |
| 6B | Ratio IHF length ventral forebrain C57 versus BTBR | n = 6 C57, n = 6 BTBR | Mann-Whitney test | 0.0152 |
| 6B | Ratio IHF length mid-horizontal forebrain C57 versus BTBR | n = 6 C57, n = 6 BTBR | Mann-Whitney test | 0.0022 |
| 6B | Ratio IHF length dorsal forebrain C57 versus BTBR | n = 6 C57, n = 6 BTBR | Mann-Whitney test | 0.0022 |
| 6E | Ratio IHF width C57 versus BTBR | n = 7 C57, n = 6 BTBR | Mann-Whitney test | 0.0012 |
| 6G | NESTIN fluorescence intensity ROSTRAL 0 – 50 µm lateral to the IHF, C57 versus BTBR | n = 7 C57, n = 6 BTBR | Mann-Whitney test | 0.4452 |
| 6G | NESTIN fluorescence intensity ROSTRAL 50 – 100 µm lateral to the IHF, C57 versus BTBR | n = 7 C57, n = 6 BTBR | Mann-Whitney test | 0.0221 |
| 6H | NESTIN fluorescence intensity CAUDAL 0 – 100 µm lateral to the IHF, C57 versus BTBR | n = 7 C57, n = 6 BTBR | Mann-Whitney test | 0.4452 |
| 6K | E14 SOX9-positive MZG, C57 versus BTBR | n = 9 C57, n = 8 BTBR | Unpaired t test | 0.0003 |
| 6K | E15 SOX9-positive MZG, C57 versus BTBR | n = 6 C57, n = 6 BTBR | Mann-Whitney test | 0.0022 |
| 6L | E15 SOX9-positive MZG 0-50 µm bin, C57 versus BTBR | n = 6 C57, n = 6 BTBR | 2-way ANOVA with Sidak's multiple comparison test | 0.0002 |
| 6L | E15 SOX9-positive MZG 50-100 µm bin, C57 versus BTBR | n = 6 C57, n = 6 BTBR | 2-way ANOVA with Sidak's multiple comparison test | <0.0001 |

| | | | | |
|---|---|---|---|---|
| 6L | E15 SOX9-positive MZG 100-150 µm bin, C57 versus BTBR | n = 6 C57, n = 6 BTBR | 2-way ANOVA with Sidak's multiple comparison test | 0.0003 |
| 6L | E15 SOX9-positive MZG 150-200 µm bin, C57 versus BTBR | n = 6 C57, n = 6 BTBR | 2-way ANOVA with Sidak's multiple comparison test | 0.0072 |
| 6L | E15 SOX9-positive MZG 200-250 µm bin, C57 versus BTBR | n = 6 C57, n = 6 BTBR | 2-way ANOVA with Sidak's multiple comparison test | 0.8128 |
| 6L | E15 SOX9-positive MZG 250-300 µm bin, C57 versus BTBR | n = 6 C57, n = 6 BTBR | 2-way ANOVA with Sidak's multiple comparison test | >0.9999 |
| 7E | Total midline length E13 C57 versus BTBR | n = 7 C57, n = 8 BTBR | Mann-Whitney test | 0.1520 |
| 7E | Total midline length E14 C57 versus BTBR | n = 9 C57, n = 8 BTBR | Unpaired t test | 0.9152 |
| 7E | Total midline length E15 C57 versus BTBR | n = 4 C57, n = 8 BTBR | Unpaired t test | 0.5507 |
| 7F | Percent EdU-positive/DAPI-positive MZG E13 | n = 6 C57, n = 8 BTBR | Mann-Whitney test | 0.9497 |
| 7F | Percent EdU-positive/DAPI-positive MZG E14 | n = 9 C57, n = 7 BTBR | Mann-Whitney test | 0.0115 |
| 7F | Percent EdU-positive/DAPI-positive MZG E15 | n = 4 C57, n = 7 BTBR | Mann-Whitney test | 0.0121 |
| 7G | Percent EdU-positive/KI67-positive MZG E13 | n = 6 C57, n = 8 BTBR | Mann-Whitney test | 0.5728 |
| 7G | Percent EdU-positive/KI67-positive MZG E14 | n = 9 C57, n = 7 BTBR | Mann-Whitney test | 0.1142 |
| 7G | Percent EdU-positive/KI67-positive MZG E15 | n = 4 C57, n = 7 BTBR | Mann-Whitney test | 0.0061 |
| 7H | Percent EdU-positive/KI67-negative MZG E13 | n = 6 C57, n = 8 BTBR | Mann-Whitney test | 0.5728 |
| 7H | Percent EdU-positive/KI67-negative MZG E14 | n = 9 C57, n = 7 BTBR | Mann-Whitney test | 0.1142 |

| | | | | |
|---|---|---|---|---|
| 7H | Percent EdU-positive/KI67-negative MZG E15 | n = 4 C57, n = 7 BTBR | Mann-Whitney test | 0.0061 |
| 7I | EdU-positive cells IHF base E13 | n = 7 C57, n = 8 BTBR | Mann-Whitney test | 0.0003 |
| 7I | EdU-positive cells IHF base E14 | n = 9 C57, n = 7 BTBR | Mann-Whitney test | 0.0086 |
| 7I | EdU-positive cells IHF base E15 | n = 4 C57, n = 7 BTBR | Mann-Whitney test | 0.1636 |
| 7J | Density EdU-positive/KI67-positive cells IHF base E13 | n = 7 C57, n = 8 BTBR | Mann-Whitney test | 0.0007 |
| 7J | Density EdU-positive/KI67-positive cells IHF base E14 | n = 9 C57, n = 7 BTBR | Mann-Whitney test | 0.7577 |
| 7J | Density EdU-positive/KI67-positive cells IHF base E15 | n = 4 C57, n = 7 BTBR | Mann-Whitney test | 0.4121 |
| 7K | Density EdU-positive/KI67-negative cells IHF base E13 | n = 7 C57, n = 8 BTBR | Mann-Whitney test | 0.3450 |
| 7K | Density EdU-positive/KI67-negative cells IHF base E14 | n = 9 C57, n = 7 BTBR | Mann-Whitney test | 0.0934 |
| 7K | Density EdU-positive/KI67-negative cells IHF base E15 | n = 4 C57, n = 7 BTBR | Mann-Whitney test | 0.0727 |
| S1D | Ratio anterior-posterior fused septum length pCCD humans | n = 9 control, n = 6 pCCD | Mann-Whitney test | 0.0044 |
| S3A | HC length chr4:rs6397070 SNP | n = 10 heterozygous, n = 21 homozygous | Kruskal-Wallis ANOVA with Dunn's multiple comparison test | > 0.9999 |
| S3B | HC length chr9:rs29890894 SNP | n = 15 heterozygous, n = 16 homozygous | Kruskal-Wallis ANOVA with Dunn's multiple comparison test | > 0.9999 |
| S3C | HC length chr15:rs31781085 SNP | n = 16 heterozygous, n = 15 homozygous | Kruskal-Wallis ANOVA with Dunn's multiple comparison test | > 0.9999 |

| | | | | |
|---|---|---|---|---|
| S3D | HC length *Draxin* mutation | n = 39 heterozygous, n = 29 homozygous | Kruskal-Wallis ANOVA with Dunn's multiple comparison test | > 0.9999 |
| S3E | E13 EdU-positive cells within base IHF C57 versus BTBR | C57 n = 7, BTBR n = 8 | Mann-Whitney test | 0.0003 |
| S3E | E14 EdU-positive cells within base IHF C57 versus BTBR | C57 n = 10, BTBR n = 6 | Mann-Whitney test | 0.0086 |
| S3E | E15 EdU-positive cells within base IHF C57 versus BTBR | C57 n = 4, BTBR n = 7 | Mann-Whitney test | 0.1636 |
| S3F | E13 volume base IHF C57 versus BTBR | C57 n = 7, BTBR n = 8 | Mann-Whitney test | 0.014 |
| S3F | E14 volume base IHF C57 versus BTBR | C57 n = 10, BTBR n = 6 | Mann-Whitney test | 0.016 |
| S3F | E15 volume base IHF C57 versus BTBR | C57 n = 4, BTBR n = 7 | Mann-Whitney test | 0.0121 |

AC = anterior commissure, CC = corpus callosum, cCCD = complete corpus callosum dysgenesis, Chr = chromosome, HC = hippocampal commissure, IHF = interhemispheric fissure, MZG = midline zipper glia, pCCD = partial corpus callosum dysgenesis, SNP = single nucleotide polymorphism.
